# BiomarkerKB: An Integrated Knowledgebase Supporting Biomarker-Centric Exploration of Biomedical Data

**DOI:** 10.64898/2026.01.26.701395

**Authors:** Daniall Masood, Mariia Kim, Jeet Vora, Robel Kahsay, Patrick McNeeley, Sean Kim, Sujeet Kulkarni, Darren A Natale, Srinivasan Ramachandran, Shakti Gupta, Mano Maurya, Cristian G Bologa, Thomas S DeNapoli, Vincent T Metzger, Praveen Kumar, Nasheath Ahmed, John Erol Evangelista, Sean C Kelly, Jorge L. Sepulveda, Avi Ma’ayan, Jonathan Silverstein, Deanne M Taylor, Daniel J Crichton, Ashish Mahabal, Jeremy J Yang, Christophe G Lambert, Shankar Subramaniam, Mike Tiemeyer, Rene Ranzinger, Raja Mazumder

**Affiliations:** Department of Biochemistry and Molecular Medicine, School of Medicine and Health Sciences, The George Washington University, Washington, DC, USA; Complex Carbohydrate Research Center, University of Georgia, Athens, Georgia, USA; Protein Information Resource, Georgetown University Medical Center, Washington, DC, USA; Department of Bioengineering, University of California, San Diego, La Jolla, California, USA; Department of Internal Medicine, School of Medicine, Translational Informatics Division, University of New Mexico, Albuquerque, New Mexico, USA; Department of Pharmacological Sciences and Artificial Intelligence and Human Health, Mount Sinai Center for Bioinformatics, Icahn School of Medicine at Mount Sinai, New York, New York, USA; Jet Propulsion Laboratory, California Institute of Technology, Pasadena, California, USA; Department of Pathology, School of Medicine and Health Sciences, The George Washington University, Washington, DC, USA; Department of Biomedical Informatics, University of Pittsburgh School of Medicine, Pittsburgh, Pennsylvania, USA; Department of Biomedical and Health Informatics, The Children’s Hospital of Philadelphia, Philadelphia, Pennsylvania, USA; Department of Pediatrics, University of Pennsylvania Perelman School of Medicine, Philadelphia, Pennsylvania, USA; California Institute of Technology, Pasadena, California, USA; School of Medicine, Department of Internal Medicine, Translational Informatics Division, University of New Mexico, Albuquerque, New Mexico, USA

## Abstract

Biomarkers are essential tools for disease detection, risk assessment, therapeutic monitoring, and precision medicine. However, biomarker data are dispersed across heterogeneous resources, inconsistently reported in the literature, and rarely standardized for computational use. This fragmentation limits reproducibility, cross-study integration, and the discovery of novel biomarker and disease relationships. We developed BiomarkerKB, a knowledgebase designed to harmonize and integrate biomarker information under a standardized data model. The model follows the FDA-NIH BEST biomarker definition and captures both core fields (biomarker entity, disease/condition, exposure agent) and contextual metadata (specimen, biomarker role, evidence, provenance). Biomarker data and related annotations were either curated from publications or collected from public resources (e.g., OpenTargets, GWAS Catalog, ClinVar, CIViC, OncoMX) and were also contributed by the Common Fund Data Coordinating Centers and the Early Detection Research Network (EDRN). Standardization was achieved using ontologies and reference resources such as Disease Ontology, UBERON, UniProtKB, and HUGO Gene Nomenclature Committee (HGNC) gene symbols. BiomarkerKB data were ingested into a Neo4j-based knowledge graph and integrated with the Common Fund Data Ecosystem (CFDE) Knowledge Graph. The initial release of BiomarkerKB contains over 200,000 biomarker-disease associations spanning genes, proteins, metabolites, glycans, and chemical elements. The knowledge graph comprises more than 300,000 nodes and 1.2 million edges, enabling structured exploration of biomarker relationships within CFDE data as demonstrated through the knowledge graph query-based use cases presented in this study. A publicly accessible web portal (https://biomarkerkb.org) provides keyword search, filtering, data downloads, and access to graph visualization to support both researchers and computational analyses. BiomarkerKB addresses a critical gap in biomarker informatics by providing an integrated, FAIR (Findable, Accessible, Interoperable, and Reusable), and unified framework for biomarker knowledge exploration and discovery.

## INTRODUCTION

Biomarkers are fundamental to modern biomedical research and clinical practice, enabling the identification of disease, prediction of therapeutic response, and monitoring of patient outcomes^1–3^. Over the past two decades, the rapid expansion of high-throughput technologies, clinical trials, and electronic health records (EHRs) has led to an unprecedented growth of biomarkers and related annotations, including genetic, proteomic, glycomic, and metabolomic measurements. Despite their central role in general healthcare and advancing precision medicine, biomarker data, including how they are analyzed, are often siloed in domain-specific databases, buried within unstructured text in publications, or inconsistently reported across clinical guidelines^4–7^. This fragmentation hampers systematic integration, reproducibility, and cross-study comparability, ultimately limiting the translational potential of biomarker discoveries.

While several databases have been developed to capture biomarker-relevant annotations, to the best of our knowledge, they primarily focus on the biomolecular entity (e.g., gene, protein) without using a standardized biomarker data model that is extensible to different types of biomarker entities. Such resources include Cancer Genome Interpreter^8^, ResMarkerDB^9^, COSMIC^10^, MarkerDB^11^, NHGRI-EBI Genome Wide Association Study catalog (GWAS)^12^, Clinical Interpretation of Variants in Cancer (CIViC)^13^, TheMarker^14^, and several disease-specific biomarker resources and publications^15–18^. While the availability of these datasets is extremely valuable, each group has its own scope and focus when reporting biomarker data. Furthermore, contextual information about how a biomarker changes in relation to a condition, such as an increase in concentration or a decrease in expression, is usually not mentioned. The absence of a unified biomarker data model that incorporates both entity definitions and contextual result changes has left a critical gap for data integration and computational analyses.

To address this challenge, we developed BiomarkerKB, a knowledgebase designed to harmonize, integrate, and contextualize biomarker data under a standardized data model. BiomarkerKB builds on the FDA-NIH Biomarker Working Group’s (FNBWG) definition of a biomarker as a “characteristic that is measured as an indicator of normal biological processes, pathogenic processes, or responses to an exposure or intervention, including therapeutic interventions”^3^. This ensures that biomarkers are represented not only as entities (e.g., gene, protein, glycan, metabolite) but also as structured relationships linking result changes to health and disease states. This framework enables the consolidation of biomarker knowledge from diverse sources.

In parallel, we constructed a Biomarker Knowledge Graph (BKG) that highlights context-specific biomarker relationships enriched with provenance, evidence, and measurement metadata. The BKG is implemented in Neo4j and integrated with the NIH Common Fund Data Ecosystem (CFDE) Data Distillery Knowledge Graph (DDKG)^19^, providing interoperability with broader biomedical datasets.

BiomarkerKB represents a step toward making biomarker data FAIR (Findable, Accessible, Interoperable, and Reusable)^20^, supporting both translational research and clinical research.

## MATERIALS AND METHODS

### Biomarker Data Model Development

The biomarker data model was developed during the preliminary phase of this project, aiming to standardize cancer and COVID-19 biomarkers^21–24^. The model follows the FNBWG definition for Biomarkers in the BEST (Biomarkers, EndpointS, and other Tools) Resource. The data types and relationships shown in Figure 1 follow this definition and incorporate contextual data deemed valuable and provide an updated view of the biomarker data model. The core fields are denoted in green and are directly related to the definition. These core fields include “biomarker”, “assessed_biomarker_entity”, “assessed_biomarker_entity_ID”, “condition”, “condition_ID”, “exposure_agent”, and “exposure_agent_ID.” The “biomarker” field follows the biomarker definition and incorporates the “assessed_biomarker_entity” preceded by the change describing it. The contextual fields, labeled in blue, provide additional information about the biomarker, such as “specimen”, “biomarker role”, and “evidence”.

**Figure 1.**
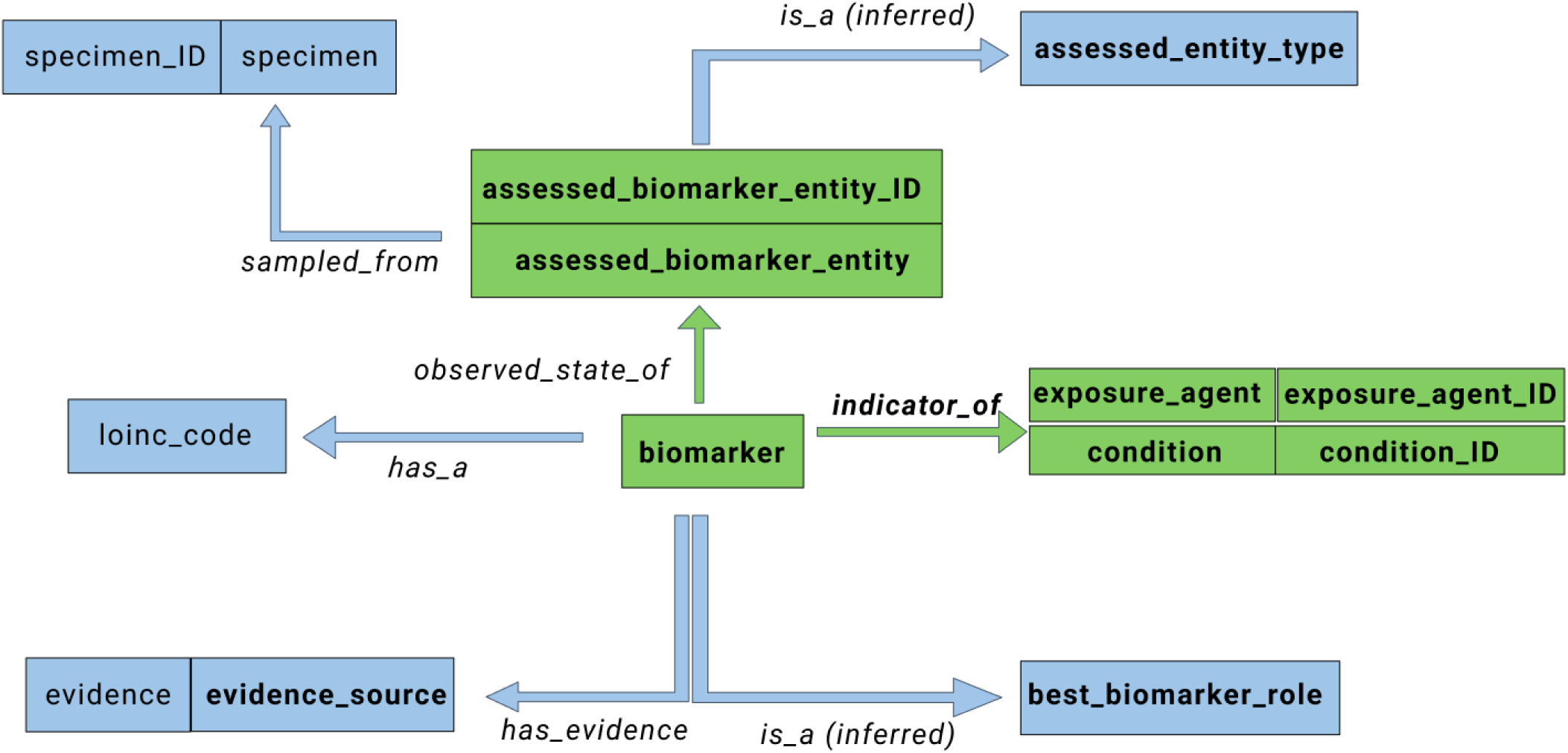
Biomarker Data Model. Schematic representation of the BiomarkerKB data model, developed by the Biomarker Partnership. Core fields (green) represent essential elements of the FDA–NIH BEST biomarker definition, including biomarker entity, condition, and exposure agent. Contextual fields (blue) capture additional metadata such as specimen, biomarker role, and evidence source, supporting extensibility and reproducibility.

### Data Collection and Curation

Identifying biomarker data that conformed to the biomarker data model was challenging because not all data sources or publications referred to or structured biomarker information in the same way. To qualify as biomarker data, the records had to represent a change within an entity and be associated with a condition or its intervention. Most sources reported biomarker data as the presence of an entity, without explicitly specifying the change associated with the condition or response to treatment.

Initial focus was on data curation from well-established resources that reported biomarker data consistent with the biomarker definition. These resources included OpenTargets^25^, the GWAS Catalog^12^, ClinVar^26^, CIViC^13^, and OncoMX^24^. This project focuses on collecting and integrating biomarker data only from resources that have permissive licenses (e.g., CC0, CC BY 4.0 that permits unrestricted commercial and non-commercial use). This also promotes FAIR principles by enabling transparent provenance tracking, reproducibility, and broad reusability of public biomarker datasets. More restrictive resources, such as COSMIC^10^, OncoKB^27^, and MarkerDB^11^ are not fully integrated within BiomarkerKB. However, biomarkers within BiomarkerKB are cross referenced to these resources whenever possible in order to provide further information and to promote research collaboration and interoperability.

The minimum data collected from the other resources comprised the model’s core fields, all of which were required for inclusion in BiomarkerKB. Where possible, other contextual data was obtained from the resource. These include specimen source (e.g., blood, saliva) and evidence source. The biomarker roles include risk, diagnostic, prognostic, monitoring, predictive, response, and safety. Risk biomarkers reflect measurable features that signal an individual’s increased likelihood of developing a disease or medical condition before any clinical signs are present. Diagnostic biomarkers are used to establish whether a disease or condition is present. Prognostic biomarkers provide information about how a disease is likely to progress over time. Monitoring biomarkers are measurements obtained serially to track changes in disease status or to evaluate ongoing biological effects related to treatment or environmental exposure. Predictive, response, and safety biomarkers are used to characterize outcomes associated with therapeutic or environmental interventions. Predictive biomarkers identify individuals who are more or less likely to benefit from, or be harmed by, a specific intervention. Response biomarkers capture measurable biological changes that occur following exposure to a treatment or environmental factor. Safety biomarkers are used to evaluate the potential for, or extent of, harmful effects resulting from an intervention or exposure^3^.

All collected data is documented with detailed descriptions of acquisition and harmonization procedures through the IEEE-standard BioCompute Objects^28^ (https://biocomputeobject.org/) and is freely accessible online (https://data.biomarkerkb.org).

#### Manual Biomarker Data Collection

Manual Biomarker Data Collection included extraction from OncoMX, UniProtKB, and the Metabolomics Workbench. OncoMX, previously developed in collaboration with EDRN^29^ and later extended to include COVID-19 biomarkers during the pandemic^21^. The minimum data collected comprised the model’s core fields, all of which were required for inclusion in BiomarkerKB, and contextual data such as specimen source and evidence source were incorporated when available. Identifiers for core and contextual entities (assessed_biomarker_entity_ID, specimen_ID, etc.) were sourced from relevant ontologies or well-established databases, including Disease Ontology (DO) and UBERON. A biomarker_role, populated with biomarker types (risk, diagnostic, prognostic, monitoring, safety, predictive, or response) defined by the FNBWG, was assigned to each biomarker. This role was either inferred by manual curators, provided directly by the resource, or inferred from the evidence. UniProtKB, a comprehensive protein knowledgebase, also contained biomarker evidence even though it primarily provides protein sequences and annotations^30^. A search for the word “biomarker” was conducted, and for each result the flat text file was downloaded for biomarker data extraction. These expert-reviewed entries provided titles of papers and curated evidence sentences, allowing mapping into “biomarker,” “assessed_biomarker_entity,” “specimen,” and “condition.” The UniProtKB accession ID was mapped into “assessed_biomarker_entity_id,” and “assessed_entity_type” was always either protein or gene. “specimen_ID” and “condition_ID” were mapped from UBERON and DO, and the “biomarker_role” was inferred from the text provided in each entry. “Evidence_source” consisted of the UniProtKB accession and associated PMIDs, and “Evidence” was populated with curated sentences from the UniProtKB/Swiss-Prot entry file. Metabolite biomarker data were obtained from the Metabolomics Workbench, focusing specifically on metabolites used within the U.S. Uniform Newborn Screening Program (https://www.hrsa.gov/advisory-committees/heritable-disorders/rusp). Relevant metabolite biomarkers were extracted from datasets and documentation describing clinically validated screening targets, with associated diagnostic metabolites identified and compiled for each disorder in the screening panel. Only metabolites directly referenced as part of established newborn screening assays were included, and annotation fields such as metabolite names, associated conditions, and clinical relevance were retained for downstream harmonization into the biomarker data model. The resulting dataset provides a structured collection of metabolite biomarkers linked to newborn-screened conditions, supporting analyses of clinically actionable metabolic biomarkers.

#### Automated Biomarker Data Collection

Automated Biomarker Data Collection was performed across multiple large-scale resources, including ClinVar, GWAS, OpenTargets, CIViC, and SenNet each of which required tailored filtering, harmonization, and mapping steps to adhere to the biomarker data model. ClinVar and GWAS were filtered for cancer-related variations using keywords such as “cancer” and “carcinoma,” followed by harmonization of disease fields to ensure unique biomarker–disease associations. Variants reported as dbSNP rsIDs were mapped as “presence of mutation in (gene)” in the biomarker field, with gene symbols mapped to assessed_biomarker_entity and rsIDs mapped to assessed_biomarker_entity_id. Cancer conditions were mapped to condition and corresponding DOIDs. Specimen values were expanded from shorthand (e.g., “Bld” to “blood”) and mapped to UBERON IDs, and Logical Observation Identifiers Names and Codes (LOINC) codes were assigned when possible through fuzzy matching of gene-variant descriptions to LOINC term descriptions. Evidence source identifiers were constructed from each database’s internal IDs, and biomarkers were inferred to be “gene biomarkers,” with best_biomarker_role assigned for ClinVar, GWAS, and OpenTargets, and mapped directly from CIViC evidence types.

OpenTargets required integration of three files, evidence, targets, and cancerBiomarkers to connect internal and ENSEMBL identifiers before extracting gene-level variant biomarkers. CIViC data were processed from the “Evidence TSV” file by mapping variants, associated genes, cancer conditions, DOIDs, evidence URLs, and variant identifiers into the biomarker model, with LOINC and specimen mappings again performed through matching of gene and variant descriptors. SenNet biomarkers focused on cell senescence biomarkers, using downloadable biomarker annotation data from the SenNet Consortium (https://sennetconsortium.org/). Because the “Biomarker” field did not describe a measurable change, its entries were mapped into assessed_biomarker_entity with corresponding identifiers mapped into assessed_biomarker_entity_id; “Context” terms such as “aging” or “obesity” were mapped into condition, with either DOID or Human Phenotype Ontology IDs used for condition_id depending on whether the context reflected a disease or phenotype. assessed_entity_type was derived from the “Biomolecule” field, specimens from the “Tissue” field, and citations were mapped to evidence_source, while the biomarker field remained blank for this dataset.

#### PubMed Central Biomarker Data Collection

A custom dataset was generated through manual curation of biomarker gene sets identified in PubMed Central^31^ via Rummagene^32^. The term “biomarker” was submitted to the Rummagene web server to obtain gene sets that match the search term. The search results produced by Rummagene were manually reviewed, and publications that explicitly associated specific conditions or exposures with biomarker gene sets were manually selected for inclusion. For each selected publication, the reported biomarker gene sets were extracted and verified by confirming the presence and relevance of each gene set within the context of the study findings. Only gene sets that were clearly supported by experimental or observational evidence in the corresponding manuscript were included in the final dataset. The resulting curated dataset captures biomarker panels and other biomarker gene sets linked to a range of conditions and exposures. The primary intended use of this collection is to support downstream analyses focused on identifying, comparing, and characterizing biomarker panels associated with human health conditions.

#### LLM-based Biomarker Data Collection

Biomarker data from the Early Detection Research Network (EDRN) were processed using large language model (LLM)-based agents designed to extract structured information from unstructured biomedical text. These agents from EDRN biomarker-related textual descriptions identified key biomarker attributes, including type, disease association, specimen source, and assay methodology, thereby reducing manual curation effort. Retrieval-Augmented Generation (RAG) pipelines were implemented and refined through prompt iteration and experimentation with open-weight models, and a MultiQueryRetriever was incorporated to improve retrieval robustness by returning the top five contextually relevant chunks per query, with overlapping token chunking used to preserve continuity (Supplementary File 1B). All components were implemented in LangChain using a 70 billion parameter Llama 3.3 model (https://www.preprints.org/manuscript/202411.0566; https://arxiv.org/abs/2407.21783). The publicly available EDRN RDF cancer dataset was then extracted and converted into a relational MySQL schema to support analysis and integration, and a custom SQL workflow integrated key fields from multiple tables, including biomarkers, organ_data, publications, and search_primary_names. To align extracted information with BiomarkerKB requirements, field-level extraction was performed. Four required fields, condition, status, role, and specimen, were generated in separate runs to maximize accuracy, and the outputs were subsequently aligned with the BiomarkerKB data model using further prompt optimization.

The LOINC database v2.78 was also used (https://loinc.org/downloads/) to extract chemical biomarker entities, by downloading the complete CSV file and filtering for rows labeled ‘CHEM’ or ‘PANEL.CHEM’ (11,224 x 40). The agentic AI framework was used to autonomously decide which tools and resources to use for searching and extracting evidence and annotations from biomedical articles, following the workflow outlined in Supplementary File 1A. Briefly, the agent receives a list of chemical LOINC codes, invokes a tool to retrieve LOINC descriptions, and passes these descriptions to an LLM to extract chemical-condition pairs, determine whether each represents a biomarker, classify the relationship (increase/decrease/present), and extract any evidence (PMID). If information is missing, the agent uses the PubMed MCP server to invoke E-utilities to search and fetch articles that mention the chemical-metabolite and biomarker cues in the title or abstract, and stores the retrieved articles in a searchable database. Updated information is iteratively passed to the LLM until at least three articles of evidence per pair are found. At this point, final biomarker information is written to a CSV file. This approach significantly increased the yield of chemical-disease pairs, biomarker relationships, biomarker roles, and supporting evidence, and is generalizable to any biomarker type.

Glycan biomarkers were also collected using LLM techniques. Publications were retrieved from PubMed using the NCBI Entrez API with a query designed to capture glycan-related biomarker literature published between 1900 and 2025. This search returned 4,787 records, from which titles, abstracts, and metadata were collected. A graph-based AI workflow implemented in LangGraph (https://arxiv.org/abs/2412.03801) coordinated a multi-agent system for biomarker extraction, beginning with relevance screening to determine whether each article contained meaningful glycobiology and disease content and whether full-text processing was required (Supplementary File 1C). For open-access publications, PMC XML files were retrieved, parsed into sentences, and tagged with section identifiers and sentence indices. Glycan entity recognition was performed using an LLM-based extraction agent that identified glycans, glycan classes, and structural motifs, followed by a normalization module to map heterogeneous glycan nomenclature to canonical forms. Longer articles had introduction and methods sections lightly summarized to reduce noise. Biomarker extraction was conducted by a dedicated agent that identified glycan-change-disease relationships and linked each biomarker to specific evidence sentences. An ontology-mapping agent attempted to align glycan terms with GlyTouCan IDs^33^, map diseases to DOIDs, map specimens to UBERON IDs and Cellosaurus IDs^34^, and harmonize change descriptors with the BiomarkerKB controlled vocabulary. A final evidence-checking agent verified that each biomarker claim was explicitly supported by evidence sentences, rewriting or removing unsupported entries. Validated outputs were formatted according to the BiomarkerKB ingestion schema for downstream integration. Collectively, these LLM workflows significantly expanded biomarker coverage by automating extraction, evidence verification, and ontology alignment across diverse textual and structured sources (Figure 8).

### Data Standardization

Data standardization occurred at the assessed_biomarker_entity, condition, and specimen levels. Disease Ontology was used to obtain condition_IDs, and UBERON for specimen_IDs. For the assessed_biomarker_entity, relevant entity ontologies were used to obtain the correct reference IDs. The mapping documentation is available on GitHub (https://github.com/clinical-biomarkers/biomarker-partnership/tree/main/mapping_data). For protein biomarkers, either a UniProtKB ID or a Protein Ontology (PRO) ID was used, while the HGNC symbol was used for genes. LOINC codes were used for biochemical entities and RefMet^35^ for metabolites. Standardization was important to create structure for the model and to ensure data would have cross-referencing and be used in machine learning methods.

### Knowledge Graph Construction

The BiomarkerKB Knowledge Graph (BKG) was constructed using a template web-based user interface that interfaces with a Neo4j database^36^. BiomarkerKB data was converted into nodes and edges and stored in two CSV files following the CFDE DDKG format^19^. These serialization CSV files are available for download with instructions on how to integrate the knowledge graph with the CFDE Data Distillery Knowledge Graph (https://ubkg.docs.xconsortia.org/basics/).

Instructions on how to download the BKG are available in the GitHub location for Biomarkers of Clinical Interest (https://github.com/clinical-biomarkers/Knowledge-Graph/). The BKG can be queried with Cypher to identify subgraphs within BiomarkerKB and integrate it with other compatible knowledgebases. This allowed BiomarkerKB data to be connected to the vast amount of knowledge graph assertions collected by the CFDE (e.g., gene expression in tissues and cell types, pathways, post translational modifications, drug perturbations followed by expression, etc.). The graph is also available for interactive browsing (https://biomarker-kg.maayanlab.cloud/). This website provides users with the ability to explore the neighborhood centered on an individual node, find connections between nodes, and explore use cases.

### Quality Control

During TSV ingestion and conversion into the BiomarkerKB JSON data model, the pipeline (https://github.com/clinical-biomarkers/format-converter) performs several layers of automated quality control (QC) to ensure structural integrity, identifier consistency, and evidence-level correctness. Raw header names are first extracted and verified against the expected BiomarkerKB data schema. When unexpected headers are identified, the Python standard library module difflib was used for nearest-neighbor matching and generating suggested corrections. For each detected mismatch, curators are prompted to confirm or reject the automated correction before parsing proceeds. Missing or unexpected columns are logged at the warning level for traceability purposes. Each row is then processed sequentially with QC checks applied at the biomarker, component, and specimen levels. Rows are grouped by canonical biomarker ID. Any first occurrence initializes a new BiomarkerEntry; subsequent rows undergo QC to determine whether they represent a new component or a valid update to an existing one using strict field-level core-equivalence tests. Resource prefixes and accessions are validated against metadata APIs using explicit type guards. Records failing type validation revert to TSV-provided fields rather than being discarded. When metadata-retrieved recommended names differ from those in the TSV, a warning is logged, ensuring transparent identification of terminological mismatches. For each validated ID, resource-specific URLs are constructed using metadata URL templates to ensure correct linkage to external ontologies or databases.

Evidence entries undergo a multi-step QC process as well to ensure correct assignment and deduplication. Evidence source definitions (e.g., "pubmed", "Pubmed", “PubMed”) are normalized using a namespace map to ensure consistent casing and naming. When evidence with identical (database, ID) pairs is encountered, the pipeline merges evidence texts, ensuring evidence lists remain minimal and non-redundant. The pipeline also performs explicit specimen-level validation. Before adding a specimen, it verifies the uniqueness of (specimen name, specimen_id, loinc_code) triples. Specimen links are constructed using resource-specific URL templates; empty or malformed templates produce empty URLs but do not break conversion. All unique evidence sources are compiled and validated. The pipeline fetches citation records for each evidence (e.g., PubMed references). Only citations that match the expected structure are accepted. When a citation is valid, a standardized Reference object (ID, type, URL) is appended to the citation to ensure downstream interoperability. Redundant citations detected across components are merged rather than duplicated.

### Biomarker ID Tracking

During TSV ingestion, each row is associated with a stable biomarker identifier (biomarker_id) that functions as the primary key for all aggregation and validation steps. If a row introduces a biomarker_id not previously seen, a new BiomarkerEntry object is instantiated. Encountering an already-registered ID triggers component-level reconciliation, ensuring that repeated IDs do not lead to duplicate entries. For rows sharing the same biomarker_id, all biologically defining fields (biomarker name, assessed entity, condition, exposure agent, specimen metadata) are inspected for consistency. All evidence, citations, and component-level information belonging to the same biomarker_id are accumulated into its single BiomarkerEntry. This guarantees that the JSON output contains one, and only one, canonical representation for each biomarker.

### History Tracking

To support reproducibility across database releases, an automated history-tracking system that records changes to each dataset between consecutive versions was implemented. For each new database release, the system identifies all data files associated with active BioCompute Objects (BCOs)^28^. Each BCO is linked to one tabular (TSV) output file, and only non-retired objects are processed. For each file, the pipeline computes basic statistics, including the number of fields, rows, and unique record identifiers.

To quantify changes between two consecutive releases, the workflow aligns each TSV file associated with an active BCO from the current release with the corresponding file from the previous release. When both files are present and in tabular format, they are parsed into field sets and record-identifier sets. The system then computes four delta metrics: (i) fields_added: fields present in the new release but absent in the previous one, (ii) fields_removed: fields present previously but absent in the new release, (iii) ids_added: newly observed record identifiers, and (iv) ids_removed: identifiers no longer present. For each TSV file, these deltas and the associated per-release metadata (file name, row count, field count, identifier count, and release date) are compiled into a structured JSON document (“pairs file”), which captures all changes between the two releases.

To provide a longitudinal view of how each TSV file evolves across all available releases, the system consolidates the per-release pair files into cumulative “track files.” For every release in which a TSV file appears, its historical entries are ordered chronologically and aggregated. For each release, the system computes the row count in the preceding release (row_count_last), the net change in row count relative to the previous release (row_count_change), and counts of added/removed fields and identifiers, derived from the pair file. The resulting track file contains a complete version history for each TSV file, including raw metrics, computed deltas, and temporal ordering of updates. Track files are regenerated for each new release and stored alongside the release’s JSON database artifacts. Both pair files and track files are stored in a standardized directory structure within the release’s JSON database directory.

### Biomarker Scoring Calculator

The biomarker scoring calculator was developed with a focus on extensibility and configurability, enabling users to adapt the scoring system to diverse research or clinical contexts without modifying the underlying source code. Key parameters include evidence-based weights such as the presence of clinical use (default 5 points), PubMed references, with 1 point for the first PMID and 0.2 points for each additional PMID up to a limit of 10, and evidence from non-PubMed sources, with 1 point for the first source and 0.1 points for each subsequent source. Additional considerations include 1 point for biomarkers with an associated LOINC code, and a penalty of -4 points for generic or non-specific conditions (e.g., “Cancer,” DOID:162). The framework will allow for on-the-fly customization of default weights by users in the future, which will influence the final biomarker score, ensuring flexibility across use cases (e.g., novelty vs strength of evidence).

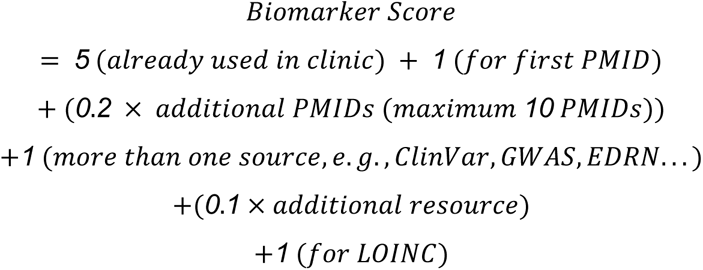

### Web Server Implementation

BiomarkerKB has separate front-end and back-end components. The back-end API component is developed using the Flask web application framework (https://flask.palletsprojects.com/en/stable/). MongoDB (https://www.mongodb.com/) is used for data storage. The front-end component is developed using the React (https://react.dev/) JavaScript library whereas Axios (https://axios-http.com/) is used to make API requests and retrieve data from the back-end layer. Docker containers (https://www.docker.com/) are used to create sandboxed environments and deploy both back-end and front-end components. An nginx (https://nginx.org/en/) web server is used to serve HTTP requests for both front-end and back-end components of the BiomarkerKB.

## RESULTS AND DISCUSSION

### Biomarker Data Model and Coverage

The development of the biomarker data model focused on enabling stable, harmonized biomarker data collection. Core biomarker data is shown in the model as the green sections (Figure 1); these are vital for defining biomarkers (biomarker, assessed_biomarker_entity, condition). Other sections of the model are considered important, but if they are not present within the paper or resource from which the biomarker is being collected, then they can be mapped from outside resources and namespaces. This data model is fashioned around the specific definition of a biomarker, which indicates that there needs to be a specific change within a molecular entity in association with a disease, treatment of a disease, or exposure. This is the central part of the biomarker data model and helps the knowledgebase remain unique and first of its kind for collecting biomarkers in a structured and harmonized fashion.

The initial release of BiomarkerKB integrates more than 200,000 biomarker-condition associations (a given biomarker can have multiple condition associations) (Table 1). Biomarkers span multiple entity types, including genes, proteins, metabolites, and glycans, reflecting the diversity of measurable entities. Cancer-related gene sequence variation biomarkers represent the largest subset, but significant coverage is also provided for metabolic, immunological, and pharmacological biomarkers through resources such as Metabolomics Workbench^37^, Library of Integrated Network-based Cellular Signatures (LINCS)^38^, and GlyGen^39^ (Figure 2).

**Figure 2.**
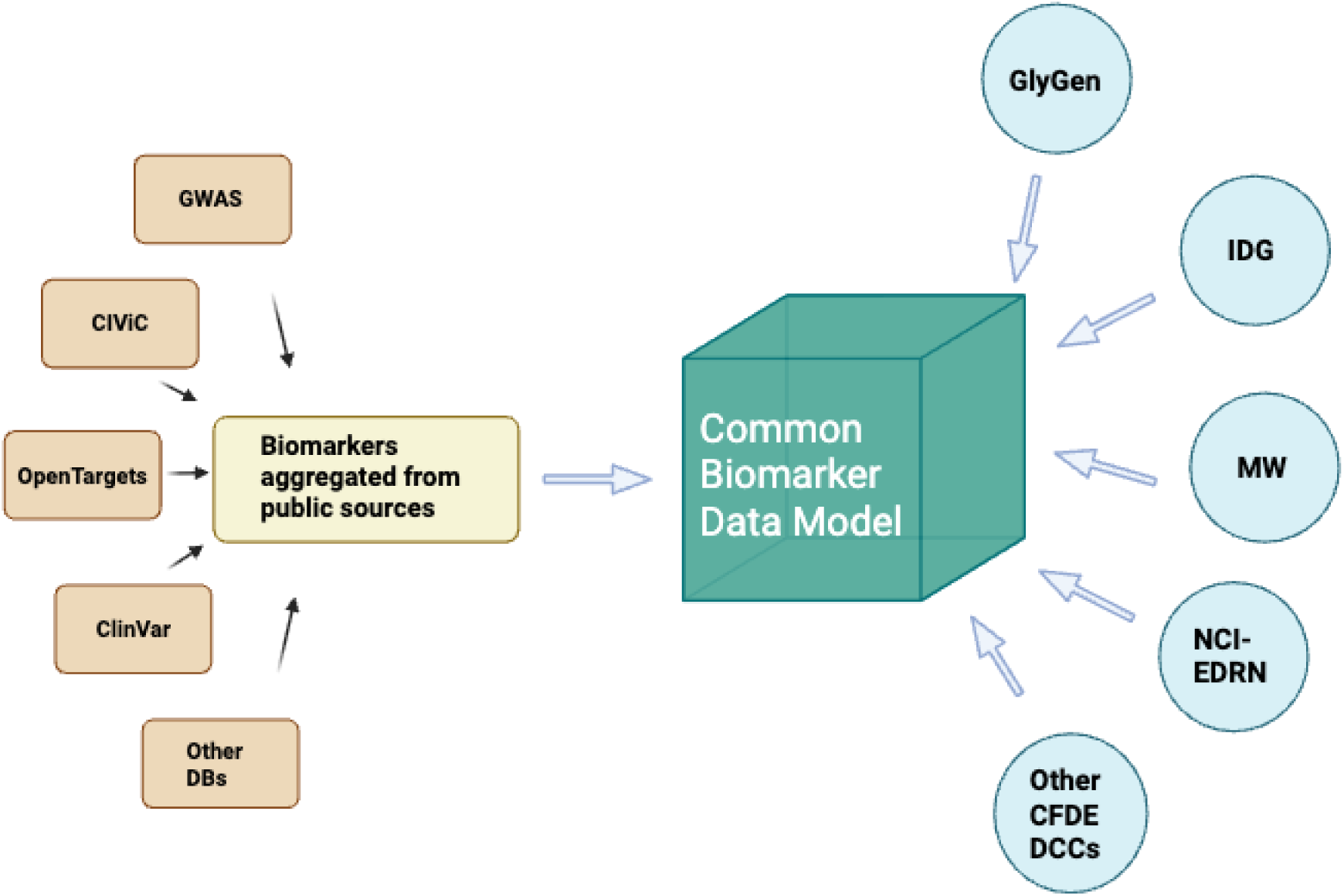
Data Integration Workflow. Overview of the data curation and integration process. Biomarker data were collected from public resources (e.g., OpenTargets, GWAS Catalog, ClinVar, CIViC, OncoMX) and contributed by CFDE Data Coordinating Centers (e.g., LINCS, Metabolomics Workbench, GlyGen, EDRN). Data were standardized using ontology references (DOID, UBERON, UniProtKB, NCBI Gene) and reformatted into the BiomarkerKB data model.

**Table 1.** Summary of Biomarker and Condition Counts in BiomarkerKB (Data Version: 2.3.1 (24/Jan/2026).

| Biomarker Data | Counts |
| --- | --- |
| Unique Biomarkers<sup>a</sup> | 205,171 |
| Biomarker-Disease Associations<sup>b</sup> | 207,314 |
| Multicomponent Biomarkers<sup>c</sup> | 118 |
| Conditions | 1166 |

<sup>a</sup>Unique biomarkers refer to the number of distinct biomarkers in the knowledge base after collapsing duplicates. These unique biomarkers are defined by the canonical biomarker ID in BiomarkerKB
<sup>b</sup>Biomarker-disease associations refer to the total number of biomarker entries within BiomarkerKB. These are defined by secondary level IDs, to help separate the unique biomarker into each instance of the biomarker associated with different conditions.
<sup>c</sup>Multicomponent biomarkers have unique canonical biomarker IDs. However, each instance of a multicomponent biomarker has multiple components within, and all share the same biomarker ID.

**Table 2.** Examples of Relationships in BiomarkerKB Knowledge Graph.

| Subject | Relation | Object | Evidence/Source |
| --- | --- | --- | --- |
| TP53 | gene_associated_with_disease<sup>a</sup> | Breast Cancer | *"TP53 mutations are frequently associated with breast cancer."*<br>(MONDO, CUI: C0006142) |
| EGFR | biomarker_type_includes_gene_product | EGFR Protein | *HGNC:3236 / GENCODE* |
| BRCA1 | is_biomarker_of | Ovarian Cancer | *ClinicalTrials.gov, NCT12345678* |
| KRAS | gene_product_plays_role_in_biological_process | Cell proliferation | *GO:0008283, GOBP ontology* |

<sup>a</sup>The relations shown in this table correspond with edge types in the knowledge graph. These edge types show direct relationships between the subject and object nodes. Some relationships within the knowledge graph are more extensive and observed over multiple edges.

Each record in BiomarkerKB is standardized according to the biomarker data model, ensuring that all entries contain the core fields (biomarker entity with its status/change, and condition or exposure agent) and associated identifiers mapped to authoritative ontologies and databases (e.g., DOID, UBERON, UniProtKB, HGNC symbol). Contextual metadata, such as biomarker role (diagnostic, prognostic, predictive, etc.) and specimen type, is captured when available.

#### Single vs. multi-component (panel) biomarkers

A biomarker may represent either a single measurable entity or a defined multi-component (panel). Most biomarkers in BiomarkerKB are single-entity biomarkers (205,053 total single entity-biomarkers), but there are currently 118 multi-component biomarkers collected and integrated into the biomarker data model and included in BiomarkerKB as well (Table 1). A multi-component biomarker consists of two or more individual biomarkers whose values are interpreted collectively, according to a specified algorithm, to indicate a biological state, disease process, or response to an exposure or intervention. The components can be different biomarker entity types (e.g., protein, gene, metabolite), and the different components can be measured by different methodologies (e.g., protein assay, genomics, etc.). Multi-component biomarkers have unique canonical biomarker IDs. All components within the multi-component biomarker share the same unique canonical ID to indicate they are part of the same biomarker and ensured they are grouped together. Both single biomarkers and panel biomarkers maintain canonical identifiers and secondary level identifiers based on disease association to support traceability, provenance, and unique reference.

### Biomarker Collection and Integration

From the integrated 205,053 single biomarkers, the breakdown of the biomarker-disease association from each resource: ClinVar contributes the largest share of biomarkers (174,602 biomarkers), followed by GWAS (5,367), CIViC (2,610), SenNet (704), OncoMX (536), OpenTargets (456), and UniProtKB (76). From EDRN 9 biomarkers have been fully integrated, and more biomarkers will be added as the EDRN resource has a total of 1500 biomarker-disease (375 of which are curated and public) associations, and from LOINC 11,124 biomarker-disease associations are available. Many biomarkers are common among the resources, especially for dbSNP biomarkers coming from GWAS, CIViC, and ClinVar. The PMC dataset provides 57 multicomponent biomarkers and a total of 1465 biomarker components and Metabolomics Workbench provides 23 biomarkers and 171 biomarker components. 122 glycan biomarkers were collected by parsing PubMed publications. UniProtKB has a low number of biomarkers extracted because biomarker-related knowledge is not an active area of curation by that group. However, some curators deemed some biomarker information important and placed this information in the comments in the flat text files. This allowed us to manually collect the 76 biomarkers from UniProtKB. Biomarker data from SenNet did not include information on change within the entity mentioned in association with disease. These biomarkers were given negative scores to help biomarkers with more relevant scores and more complete data to appear at the top of the results. However, this data is still valuable in providing further insight and will allow for further research in cell senescence to be understood.

BRCA1, BRCA2, and cancer-related conditions dominate BiomarkerKB because cancer biomarkers are well represented in structured resources such as ClinVar and GWAS, allowing us to fully develop and refine the biomarker data model. In contrast, biomarkers for non-cancer diseases remain far less organized and are often available only as unstructured free text. Having largely exhausted the collection of well-structured cancer biomarkers, our focus has shifted toward methods capable of extracting and normalizing these poorly structured biomarker descriptions, motivating us to use LLM-based approaches described below.

Although established natural language processing libraries such as SpaCy^40^, SciSpaCy^41^, and Stanza^42^ are widely used, LLMs are increasingly emerging as the preferred approach for capturing complex associations. LLMs are can extract complex relationships from the natural language found in the descriptions, as they provide better embedding (high-dimensional space, semantically rich and refined word order with positional encoding), self-attention (context-aware embeddings), and multi-headed attention (multi-perspective embeddings) of tokens (descriptions), resulting in better performance. In the first iteration, a series of Python scripts were used to preprocess (data wrangling and web scraping), process (chat template and output parsing), and post-process (data wrangling, expanding multiple specimens into individual rows, harmonizing terms to fully specified names, ontology encoding, etc.). This pipeline enabled large-scale, consistent extraction and normalization of LOINC-based biomarker tests for integration into the BiomarkerKB data model.

However, the above approach yielded sparse evidence for the biomarker and its provenance, as several tests lacked references. Similarly, biomarker status was not mentioned explicitly for several tests. This is the reason why, for the three different resources where LLM was applied, different methods had to be used for the specific use cases. Agentic AI was most useful to help extract biomarker information from free text within the LOINC database. LangChain and Llama were employed to extract biomarker information from EDRN biomarker free text. The LangGraph method was employed to collect glycan biomarkers from abstracts and full texts.

### Cross-references

BiomarkerKB enhances and contextualizes biomarker entities by cross-referencing identifiers, ontologies, and annotations from a diverse set of biomedical knowledge sources. External resources incorporated into the system include variant- and mutation-oriented databases (BioMuta^43^, Single Nucleotide Polymorphism Database (dbSNP)^44^); molecular and chemical ontologies (Cell Ontology^45^, Chemical Entities of Biological Interest (ChEBI)^46^); glycomics resources (GlyGen, GlyTouCan); gene-and expression-level repositories (Genotype-Tissue Expression (GTEx)^47^, Illuminating the Druggable Genome (IDG)^48^, Reactome^49^); clinical and laboratory terminologies (LOINC)); metabolomics and small-molecule reference databases (Metabolomics Workbench (MW), PubChem^50^); and curated biomedical knowledgebases such as UniProtKB^30^. These cross-references allow exploration of related namespaces and annotations across molecular, clinical, and phenotypic biomarker dimensions.

### Controlled Vocabulary for Biomarker Entity Types

To enable consistent annotation, interoperability, and downstream computational analysis of biomarker data, the Biomarker Entity Types Controlled Vocabulary (Supplementary Files 2-5) was developed as part of BiomarkerKB framework. The absence of standardized, controlled terminology describing the nature of biomarker entity changes, particularly across heterogeneous entities such as proteins, metabolites, glycans, and lipids, has limited data integration.

This vocabulary defines a fixed set of biomarker entity types, sequence variation, gene, protein, metabolite, glycan, DNA, RNA, cell, lipid, and image, each represented by a stable identifier (e.g., BM-0002 for protein). These entity types serve as the fundamental building blocks of BiomarkerKB’s data model, providing a consistent schema to describe the biological substance or measurable entity associated with a given biomarker. Each entity type entry follows a standardized format composed of specific line codes that capture definitions, hierarchical relationships, usage notes, and external ontology mappings. This structure allows programmatic parsing and automated linkage to external databases, ensuring that the vocabulary remains interoperable within the larger Common Fund Data Ecosystem^51^ and compatible with Biolink Model^52^ semantics.

The vocabulary was constructed through a combination of manual curation and ontology alignment. Definitions and usage notes were adapted from established bioinformatics standards, including the Sequence Ontology (SO), Chemical Entities of Biological Interest (ChEBI), Protein Ontology (PR), and the Cell Ontology (CL). Each entry was reviewed to ensure conceptual consistency and hierarchical integrity, with additional notes for mixed-composition entities (e.g., glycoproteins classified under both protein and glycan). Representative examples and canonical identifiers from BiomarkerKB (e.g., Interleukin-6, Urea, miRNA-21) were provided to illustrate correct usage and to facilitate training of automated entity recognition tools.

BiomarkerKB’s data ingestion pipeline maps all curated records to this controlled vocabulary during the normalization stage. Each biomarker is represented as a combination of a measurable descriptor (e.g., increase, presence), one of the controlled entity types (e.g., protein, metabolite), and aspect type (e.g., level, expression), and/or a modification term (e.g., glycosylation, phosphorylation). This structured format enables consistency across sources, including those derived from literature, public biomarker databases, and clinical datasets. By anchoring all biomarker entities to this standardized vocabulary, BiomarkerKB ensures that the resulting knowledgebase is semantically uniform and computationally ready for downstream applications, such as graph-based learning, embedding generation, and predictive modeling.

The controlled vocabulary for biomarker entities provides standard terms for biomarker entities that are found within BiomarkerKB (Supplementary File 2). This document lists the biomarker entity types used in the BiomarkerKB curation system. The controlled vocabulary for reporting terms lists the controlled reporting terms used to describe biomarkers in the BiomarkerKB curation system (Supplementary File 3). These nouns are used in the “biomarker” field. The controlled vocabulary for aspects lists controlled terms that represent measurable or definable aspects of a biomarker entity (Supplementary File 4). These terms indicate what property, modification, or characteristic of the biomarker is being evaluated. The controlled vocabulary for modifications lists controlled modification terms that describe chemical, structural, or enzymatic alterations to biomarker entities (Supplementary File 5). These terms are used to represent states such as methylation, phosphorylation, or glycosylation, enabling consistent annotation of modified biomarker forms.

From a computational perspective, the implementation of this controlled vocabulary transforms BiomarkerKB from a collection of heterogeneous annotations into a harmonized, machine-readable resource. Furthermore, it supports cross-resource alignment with complementary data models (e.g., OMOP, LOINC, Biolink) and facilitates reproducibility in biomarker discovery workflows. The vocabulary thus serves as a critical infrastructure component that bridges curation-level data representation and computational analysis, ensuring that biomarker data is both FAIR-compliant and ready for large-scale bioinformatics and machine learning applications.

### Knowledge Graph Construction and Features

The curated biomarker data were used to construct the BKG, implemented in Neo4j, and integrated with the CFDE Data Distillery Knowledge Graph. The graph represents biomarkers as nodes linked to conditions through using the Biolink data model. The Biolink data model provides structured relationships to describe associations between different entities (e.g. associated_with, expressed_in, related_to). This design enables queries that go beyond simple co-occurrence to capture how biomarkers change in relation to specific disease states, treatments, or exposures.

The graph currently includes over 300,000 nodes and 1.2 million edges, reflecting both curated biomarker associations and cross-links to CFDE resources. For example, gene-based biomarkers are connected to transcriptomic and pathway datasets, while metabolite biomarkers are linked to experimental assay results. These connections allow researchers to explore biomarker relationships in the context of broader molecular networks and systems biology. The graph structure also provides a substrate for computational approaches such as graph-based learning, clustering, and embedding methods to predict novel biomarker-condition associations.

Users can find a biomarker-only knowledge graph. This knowledge graph (https://biomarker-kg.maayanlab.cloud/) was developed to help users observe connections and relationships within the BiomarkerKB data^36^. This knowledge graph expands and shows visually how some biomarkers are associated with multiple conditions and how some conditions are associated with multiple biomarkers. The interface allows users to start with a node of interest and expand from there to find information and relationships that may not be observed explicitly from the tables in the knowledge base.

#### Use Case 1: Increased Interleukin-6 biomarker example

This biomarker use case highlights the ability to explore the relationship between biomarker-disease and find related treatments for the disease (Figure 3 and Supplementary File 6). In this example, increased interleukin-6, a diagnostic biomarker in breast carcinoma, is explored. The associated biomarker_id is AN6278-4. This is important in building the query as it is the starting node for the subgraph created from the query. The biomarker_id node is connected to the UNIPROTKB:P05231 node by the relationship indicated_by_above_normal_level_of. This follows the biomarker definition and builds the biomarker in the graph view. The biomarker_id node then connects to a Concept node with the preferred term “Malignant neoplasm of breast.” This then connects to another Concept node that indicates “Breast Carcinoma.” This node defines the general name of the condition and connects to three PubChem nodes with different IDs. These nodes connect to three different drug compounds, which are commonly used in treating cancer/breast cancer. These drugs were doxorubicin, paclitaxel, and everolimus. These drugs are involved in the treatment of breast cancer^53–55^. Doxorubicin is also indicated to have a wide spectrum within tumor treatment^56^. Paclitaxel has also been reviewed in cases of treating other cancer types^55^. Everolimus has been used to treat metastatic renal cancer and neuroendocrine tumors^57^. This use case demonstrates that the BiomarkerKB knowledge graph can correctly identify biomarker-disease-treatment associations.

**Figure 3.**
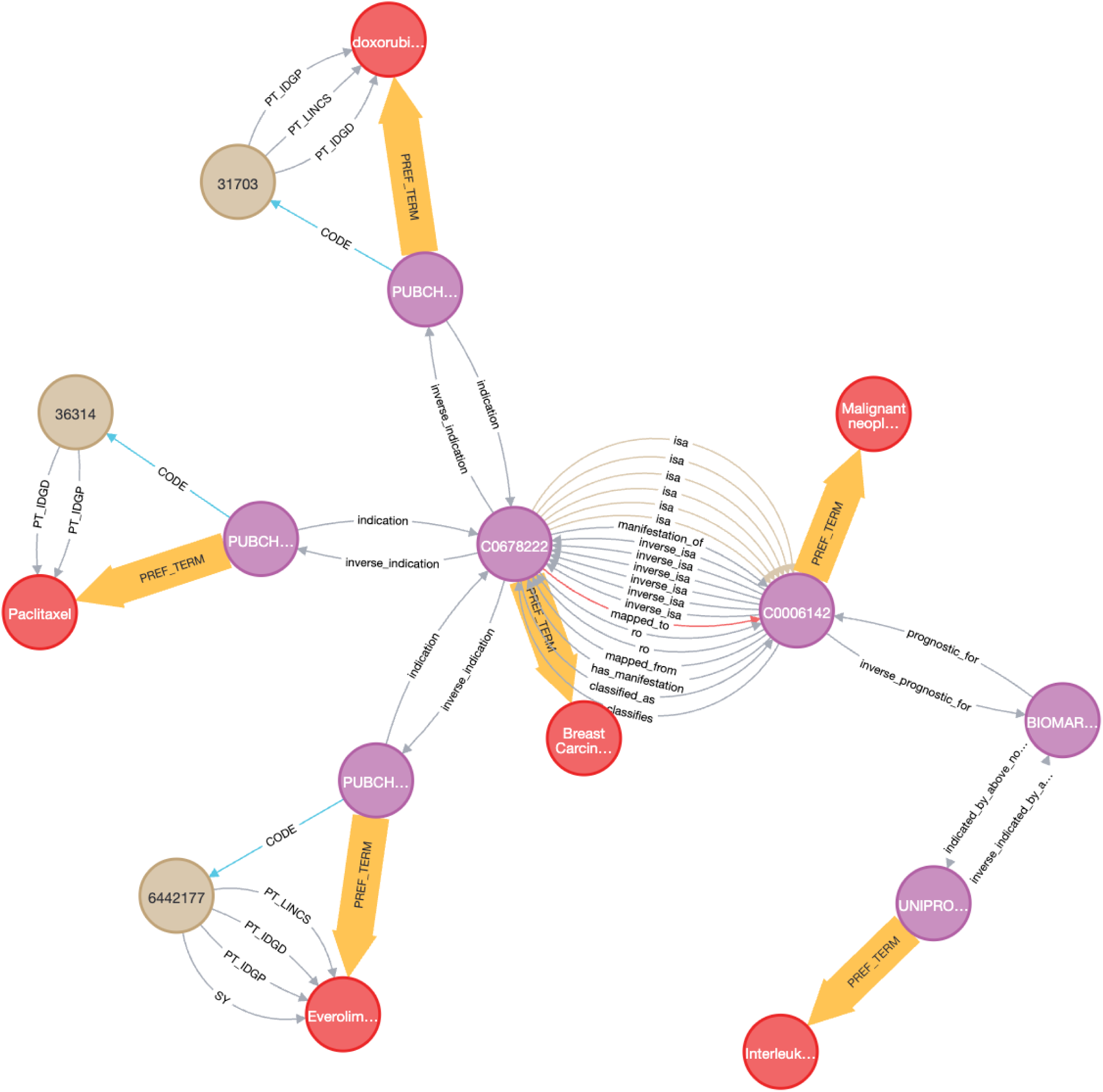
Increased Interleukin-6 biomarker knowledge graph construction. Visualization of the BiomarkerKB knowledge graph in Neo4j. Biomarkers are represented as nodes connected to conditions through context-specific is_biomarker_of relationships. An example subgraph highlights biomarker associations in breast cancer, showing links between genes, proteins, and clinical outcomes. This example of the BiomarkerKB knowledge graph focuses on interleukin-6 and breast carcinoma. Purple nodes represent standardized Concept nodes (e.g., diseases, biomarker concepts, proteins, and chemical entities), which are internal identifier nodes. Gray nodes represent source-specific identifier nodes (e.g., PubChem compound identifiers in this example). The relationships and nodes show the connection between biomarker and disease, and also identify treatments used in breast carcinoma (doxorubicin, paclitaxel, everolimus)

Because IL6 is measured in accordance to many different diseases and can be used a biomarker for different conditions, it is best the query in this use case is modeled as an example query. This example query can be expanded upon to find other relationships and information that are within the knowledge graph. This example query can be expanded through the doxorubicin treatment concept node to another disease (lymphoma) that is indicated by increased interleukin-6, and investigate if doxorubicin is a treatment. The query will still start from the BIOMARKER:AN6278-4 node, and this time shows a relationship between the increased interleukin-6 levels and lymphoma. The doxorubicin Concept node then connects to another node through the relationship of “indication” and this node connects to associated_morphology_of to another higher concept node that indicates lymphoma, which branches out to multiple concept nodes that are related to diagnosis or measuring prognosis of the disease. These nodes include several methods of biopsy. Doxorubicin is also used to treat aggressive types of lymphoma^58^.

#### Use Case 2: Exploring Cross-Disease Therapeutic Opportunities Through Biomarker-Centric Network Analysis

This biomarker use case demonstrates how the BiomarkerKB knowledge graph can be used to explore entity-disease associations beyond a single disease context and identify broader therapeutic relevance through shared molecular targets (Figure 4 and Supplementary File 7). Starting from breast cancer-associated disease concepts (Mammary Neoplasms), the query identifies genes that are recurrently linked to breast cancer and are also targeted by approved drugs. Highly druggable genes such as ERBB2, ALK, BRAF, IDH1, MTOR, and FGFR2 emerge as central nodes, each supported by multiple disease associations and varying numbers of drug interactions. ERBB2, for example, is associated with 14 distinct drugs and shows overlap with a wide range of malignancies, including lung adenocarcinoma, gastric adenocarcinoma, colorectal carcinoma, ovarian carcinoma, and glioblastoma, reflecting its well-established role as a pan-cancer oncogenic driver (Supplementary File 8). Similarly, ALK and BRAF connect breast cancer to multiple solid tumors and hematologic malignancies, while IDH1 and MTOR extend the landscape to include brain tumors, leukemias, and neurodevelopmental syndromes. These overlapping disease connections arise through gene-disease association edges and highlight how shared molecular mechanisms link multiple cancers. By integrating drug-target relationships, the graph further reveals that many therapies developed for one cancer type may have relevance in others through common biomarker pathways. Overall, this use case illustrates how BiomarkerKB enables systematic exploration of biomarker-centric disease overlap and therapeutic repurposing opportunities.

**Figure 4.**
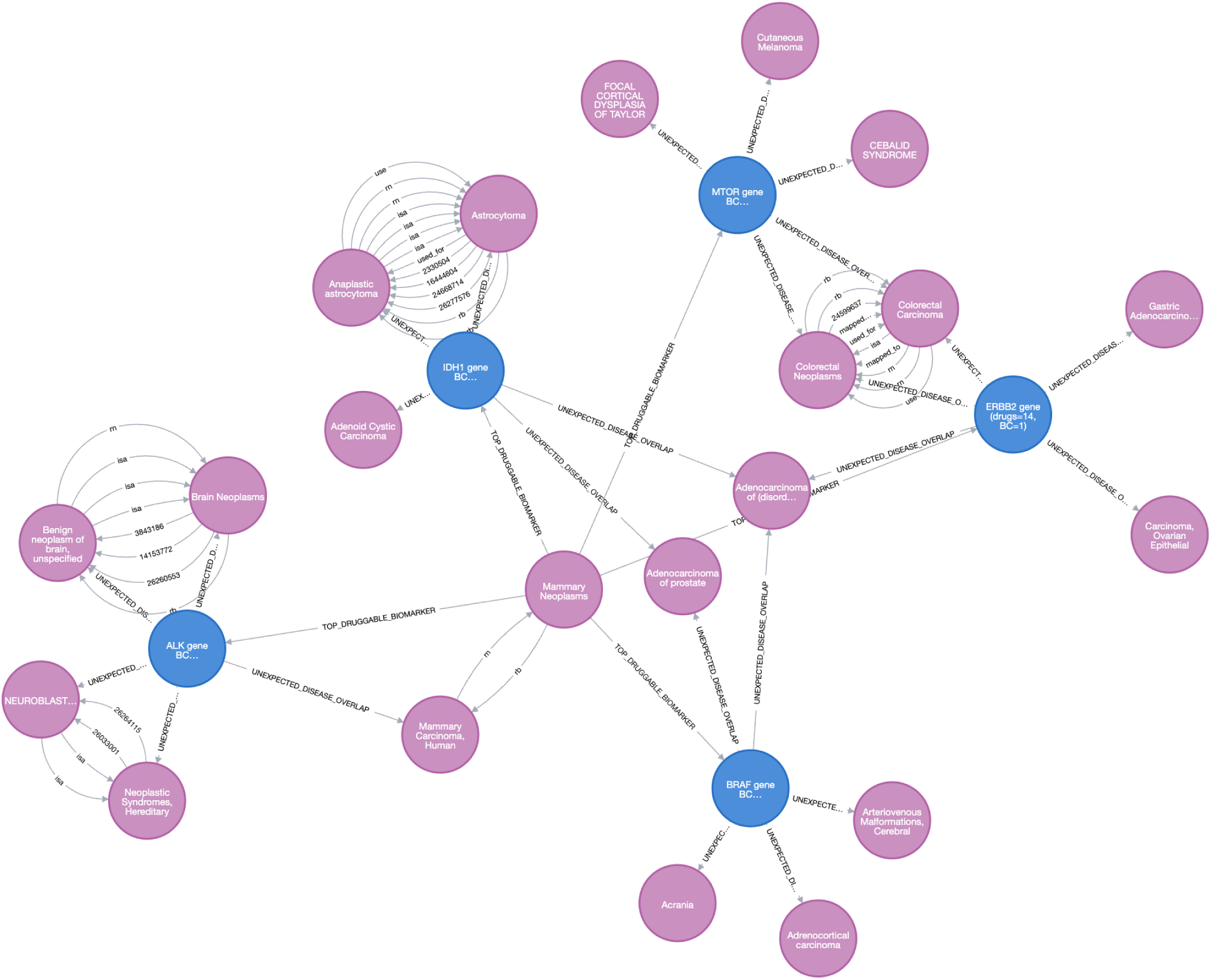
Biomarker-centric network illustrating cross-disease and drug associations derived from breast cancer-associated genes. Visualization of subgraph extracted from the BiomarkerKB knowledge graph, centered on genes associated with breast cancer and their connected diseases and drug targets. Breast cancer-associated disease concepts (Mammary Neoplasms) were used as the starting point to identify recurrently associated genes (e.g., ERBB2, ALK, BRAF, IDH1, MTOR, and FGFR2). These genes are connected to drugs via gene-drug target relationships and to additional diseases through gene-disease association edges. Peripheral disease nodes illustrate overlapping disease contexts outside the breast cancer hierarchy, highlighting shared molecular mechanisms and potential therapeutic repurposing opportunities.

#### Use Case 3: Glycosyltransferase and Tankyrase Inhibitor

This use case focuses on using the knowledge graph to find novel biomarker relationship that are not immediately evident. The query (Supplementary File 9) used to build this subgraph was focused on the research of glycosyltransferases related to cancer genome analysis^59^. The query searched through the knowledge graph to find any information that centered around the term “glycosyltransferase.” The query also focused on nodes that were related to pharmacology and treatment. Glycosylation in cancer can be a potential target for treatment because these modifications can be detected with differential expression. Due to this, the query was constrained around glycosylation (Figure 5).

**Figure 5.**
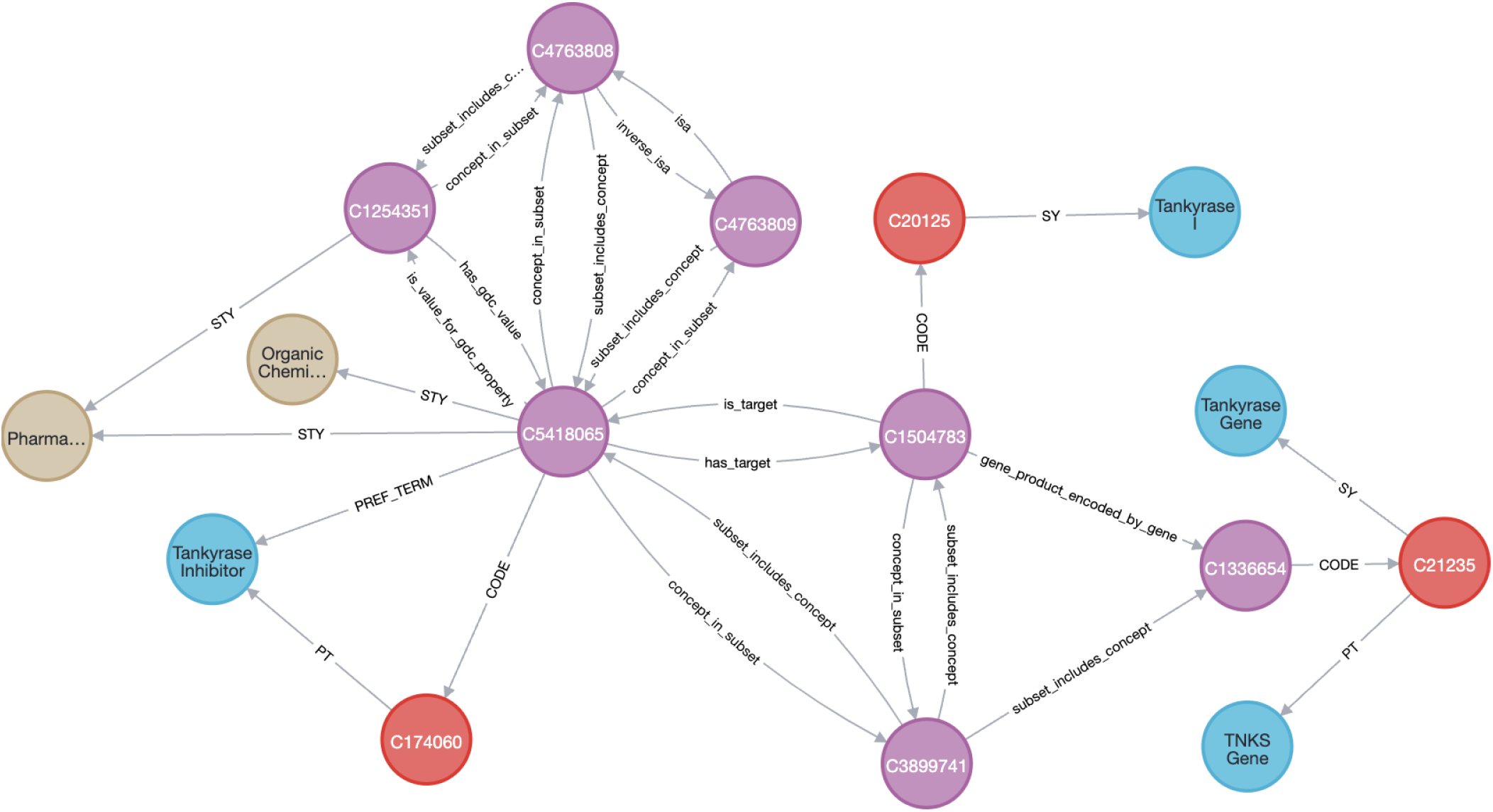
Tankyrase Inhibitor Use Case. This subgraph investigates the potential to find new biomarkers for the knowledgebase. This use case examines nodes that are related to glycosyltransferase. The node related to a tankyrase inhibitor was found and connected to the tankyrase I gene. These nodes were further described with pharmacological descriptors, indicating that this was a potential target for treatment. Purple nodes represent standardized Concept nodes (e.g., diseases, biomarker concepts, proteins, and chemical entities), which are internal identifier nodes. The CUI nodes are as follows: C1504783 = TNKS Protein, human, C3899741 = CTRP Terminology, C5418065 = Basroparib, C4763808 = GDC Terminology, C4763809 = GDC Therapeutic Agent Terminology. Other CUI Nodes have labels attached to them. This demonstrates the potential for biomarker data to be found in the graph as well as biomarker annotations.

One specific node that was found was a Concept Unique Identifier (CUI) node that had the preferred term “Tankyrase Inhibitor.” This node had a relationship “has_target” to a CUI node that had preferred terms “Tankyrase Gene”, “TKNS”, and “Tankyrase I.” This shows that the tankyrase inhibitor is being used as a treatment that targets the tankyrase gene. Tankyrase is involved in the proliferation of ovarian cancer by promoting aerobic glycolysis^60^. Although glycolysis and glycosyltransferase activity are not directly linked at the enzymatic level, the BKG suggests an indirect metabolic relationship in which glycolysis, as a catabolic pathway, can supply sugar-phosphate intermediates that serve as precursors for anabolic glycosyltransferase-mediated biosynthetic processes. However, the subgraph was not able to expand further or resolve condition- or treatment-specific relationships for this gene, indicating missing connecting data and highlighting BKG’s value as a hypothesis-generating framework for experimental investigation.

This query result exemplifies the potential for being able to find new biomarkers by finding potential targets and genes for conditions and treatments. While this example did not find fully-defined examples of biomarkers (that is, with all core information), it nonetheless demonstrated the ability to find intriguing associations for further research, including finding annotation data that can be used to add contextual information. In the future, adding more data and biomarkers to the knowledge graph will help to find new biomarkers by doing targeted queries.

#### Interactive Web Portal Functionality

The BiomarkerKB web portal (https://biomarkerkb.org) provides an interactive interface for exploring the knowledgebase (Figure 6). Users can search biomarkers by biomarker name, condition, or biomarker role, and apply filters to refine results. Each biomarker entry is linked to standardized identifiers, evidence sources, and provenance metadata.

**Figure 6.**
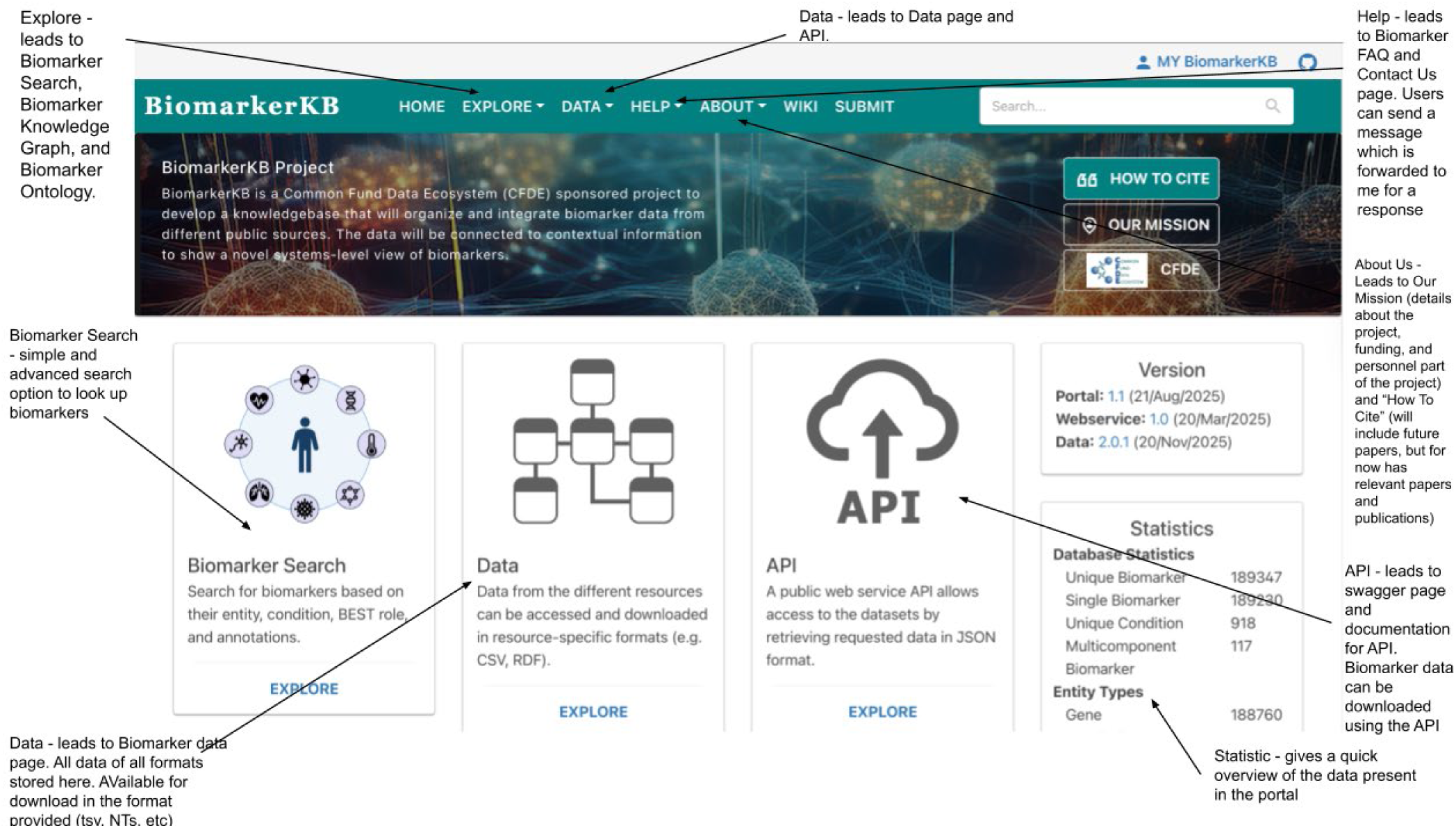
Web Portal Interface. Annotated screenshots of the BiomarkerKB web portal (https://biomarkerkb.org), illustrating search, filtering, and graph visualization capabilities.

**Figure 7.**
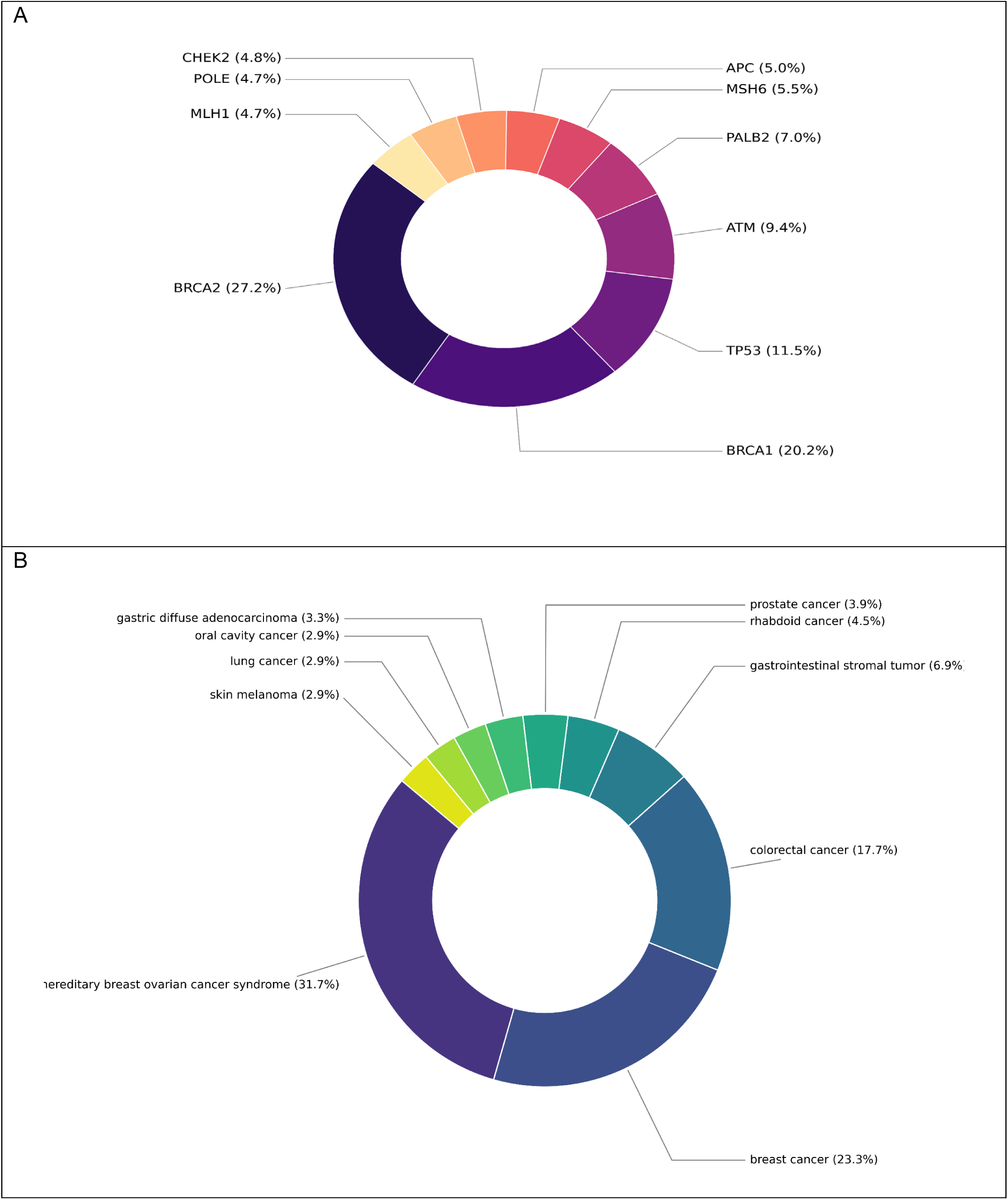
Distribution of top biomarker entities and conditions represented in the dataset. (A) Donut chart showing the top 10 assessed biomarker entities, with percentages calculated relative to the total number of biomarkers assessed. BRCA1 and BRCA2 represent the largest proportion of biomarker entities. (B) Donut chart showing the distribution of conditions with the highest number of associated biomarkers. Hereditary breast and ovarian cancer syndrome, breast cancer, and colorectal cancer account for the largest shares. Together, these visualizations highlight the predominance of cancer-related biomarkers and conditions within the dataset.

**Figure 8.**
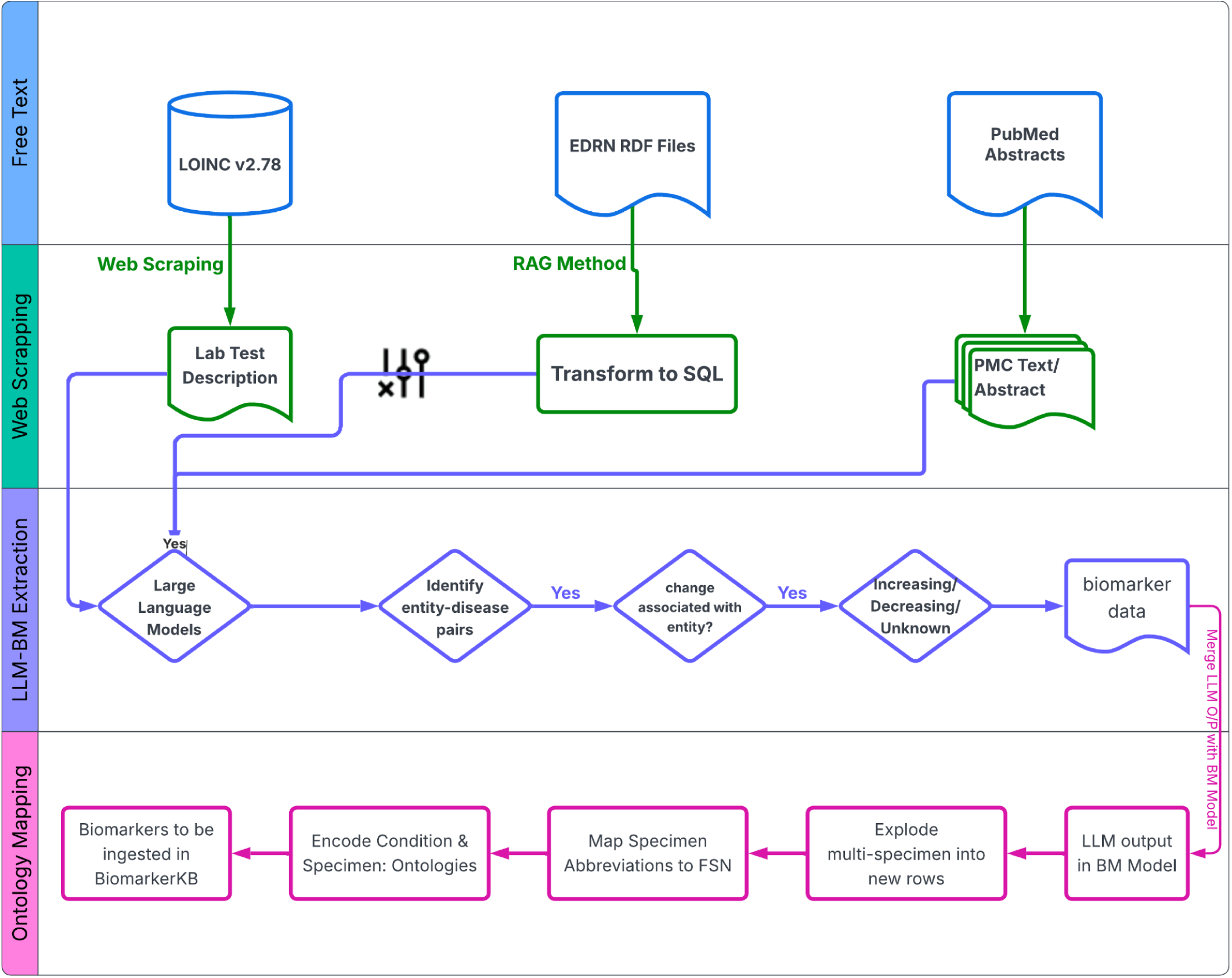
Biomarker Extraction Using Large Language Models. Different LLMs were used to extract biomarker data from different resources. LOINC, EDRN, and PubMed all have biomarker data available in unstructured free text. LLMs were used to target the free text in these different resources to extract biomarker information, map ontology IDs, and impute missing data to collect biomarkers into the Biomarker Data Model. Abbreviations in the figure: SQL= Structured Query Language; RDF= Resource Description Framework; FSN= Fully Specified Name.

Interactive graph visualizations at https://biomarker-kg.maayanlab.cloud/ enable exploration of biomarker-condition networks, while tabular views of search results at https://biomarkerkb.org facilitate bulk analysis and comparison. Data can also be downloaded in multiple formats, including CSV and JSON from data.biomarker.kb.org, to support integration with external workflows. BioCompute Objects are available for all datasets as well to document workflows, data integrity and data provenance. The system is optimized for both researchers seeking individual biomarker details and data scientists performing large-scale analyses.

Biomarker search allows for users to search for biomarkers on different criteria. Specific biomarkers can be searched for based on the actual biomarker name as well as the assessed biomarker entity to find the correct biomarker. Specific diseases can also be put into the search, and the resulting biomarkers for the diseases will be displayed. Users can also use the advanced search to specify further what biomarker they are researching. Users can input the desired biomarker entity, entity type, specimen, and disease to find the biomarker they are interested in. The advanced search also lets users further restrict parameters by ontology IDs (e.g., DOID, UBERON ID). These search options and criteria are helpful in allowing users and researchers to find the biomarker of interest efficiently.

#### AI-Assisted Query

BiomarkerKB features natural-language search through its AI-Assisted Query module that combines dense vector retrieval with large-language-model (LLM)-based semantic reasoning. The system is implemented as a Retrieval-Augmented Generation (RAG) pipeline that indexes structured biomarker records and returns context-aware answers grounded in primary database content.

Each biomarker entry is first transformed into a compact textual representation that includes its canonical identifier, assessed entities, specimen information, associated conditions, roles, and aggregated evidence excerpts. These documents are embedded using OpenAI embedding models and stored in a ChromaDB vector database. The resulting vector store supports efficient nearest-neighbor search over thousands of heterogeneous biomarker descriptions. At query time, user questions, posed in natural language, are encoded into the same embedding space, and the system retrieves the top-k most semantically relevant records. Retrieved documents are then passed, along with the query, to an LLM-based reasoning layer (via LlamaIndex) that generates a synthesized, citation-linked response. This architecture allows the system to map unconstrained biomedical questions (e.g., *“Can you show me protein biomarkers related to prostate cancer?”*) to the most relevant entries, while ensuring that the final answer is grounded in the underlying indexed data rather than generated purely from the language model.

The AI-Assisted Query feature thus provides a flexible, semantically rich interface to BiomarkerKB, supporting exploratory analysis, hypothesis generation, and rapid retrieval of biomarker-condition relationships in a way that complements traditional structured queries.

#### Data portal

The datasets processed through the BiomarkerKB data processing workflow are made available on https://data.biomarkerkb.org, providing free access to all BiomarkerKB data. The data portal enables users to search and download processed datasets by data source; these datasets also serve as inputs for generating JSON objects and powering the APIs and frontend components. Each dataset is assigned a stable BiomarkerKB dataset identifier, and a corresponding BCO is created to provide detailed documentation of the data processing workflow. The BCO is available in both machine-readable JSON format and a human-readable text format, allowing granular tracking of dataset metadata, including provenance, versioning, format, license, and attribution.

The portal interface includes links to *Home*, *FAQ*, *Release History*, *About*, *Portal*, and *API*, along with version information for the application and data. Below this section is a search bar for querying datasets. A filter panel on the left allows users to refine datasets by several categories, including Species, File Type, Molecule, Scope, and Status. The main section displays datasets in a tabular format with the following columns: *BCO ID* (identifier for the associated BioCompute Object), *Description* (summary of dataset generation or processing), *File Name* (structured dataset file name), *BCO Title (dataset name)*, and *Details* (a link to “view details” for each dataset). Selecting the “view details” link opens a detailed dataset view organized into four tabs. The All Records tab displays the dataset’s individual entries; BCO JSON provides the BCO in JSON format; README presents the BCO in text format; and Downloads includes links to download both the dataset and its BCO. The details page also includes a navigable version history, dataset tracking information, and a brief dataset description to help users understand its content and potential applications.

Overall, the data portal provides a gateway for users who wish to work with processed BiomarkerKB datasets without using the full portal or API, thereby supporting reproducible research, transparent reporting, data provenance, and efficient data sharing.

#### API Access

The BiomarkerKB backend provides a RESTful API access (https://api.biomarkerkb.org), which enables programmatic access to biomarker data and metadata. The API supports several namespaces, including `biomarker`, `auth`, `pages`, `data`, and `event`. Endpoints allow users to perform operations such as biomarker search (`/biomarker/search`), retrieval of detailed biomarker records (`/biomarker/detail/biomarker_id`), AI-assisted search (`/biomarker/ai_search`), and data download (`/data/list_download`). All responses are returned in JSON format for seamless integration with downstream tools and workflows. The API does not require authentication for public queries, facilitating open and reproducible access to biomarker information.

#### Wiki portal

The BiomarkerKB public wiki pages (https:/wiki.biomarkerkb.org) provide documentation about the project including help documentation. This is designed to update documents rapidly and provide details on many aspects of the project and maintenance of the BiomarkerKB resource. The wiki page documents data releases, version history, controlled vocabularies histories and updates, data submission and upload documentation, quality control and assessment documentation, knowledge graph documentation, portal releases, resource integration, and frequently asked questions. It provides guidance on data standards, insight into resources from where biomarker data is collected, quality control practices, and best practices for contributing new data. Contributors, users, and developers may all use the BiomarkerKB wiki to help increase understanding about the structure, methodology, and functionality of the knowledgebase. The availability of all these materials ensures transparency and consistent implementation of the BiomarkerKB framework and workflow.

#### Significance, Future Direction & Conclusion

BiomarkerKB addresses a critical gap in biomedical informatics by providing a unified, ontology-driven framework for representing biomarker data. Existing biomarker databases often limit coverage to specific entity types or lack contextual metadata about how biomarker entities change in relation to disease. In contrast, BiomarkerKB integrates heterogeneous biomarker data, standardizes representation, and contextualizes biomarker-condition relationships, thereby enabling both reproducible and scalable data integration and computational discovery. The resource allows users to search, filter, and find biomarker data based on scientific interest. Systematic harmonization of biomarker data at this scale has not been available previously, and the heterogeneous representation of biomarkers across resources has historically made integrative biomarker research difficult. Collecting biomarkers using a structured data model harmonizes the information and thus facilitates biomarker research, activities that will become even more successful as more data are added. Cross-referencing to external resources is ongoing, with the goal of reciprocal crosslinks. These will further enhance data exchange and resource findability.

The integration of BiomarkerKB into the DDKG enhances its interoperability and broadens its utility. By linking biomarkers to diverse molecular and clinical datasets, researchers can perform multi-modal analyses that were previously difficult to achieve. For example, cancer biomarkers from EDRN can be explored alongside transcriptomic perturbation data from LINCS or metabolomic signatures from Metabolomics Workbench, providing new opportunities for hypothesis generation. The knowledge graph also allows for relationships within the data to be observed and used to access novel information. Knowledge graphs can be queried and allow users to find information that previously may not have been easily accessible. The CFDE DDKG and BKG have data and information that have been well curated, so the data can be used with a high degree of reliability.

The harmonized and integrated foundation of BiomarkerKB also makes large-scale biomarker data amenable to downstream extensions, including the incorporation of aggregated EHR-derived observations to characterize biomarker distributions in healthy populations. Although defining “normal” biomarker ranges is inherently challenging due to assay heterogeneity, biological variability, and latent pathology, systematic mapping and unit harmonization represent a necessary first step that BiomarkerKB enables. The troponin case study illustrates this approach: by aggregating harmonized measurements and using median-centered summaries with box-plot visualizations, population-level trends (Supplementary File 10) can be assessed and compared against published reference patterns. Deviations from expected distributions highlight areas requiring deeper methodological investigation. Together, these analyses demonstrate how structured annotation and statistical summarization of EHR-derived biomarker data can enhance interpretability while laying the groundwork for scalable, data-driven refinement of biomarker knowledge.

Although BiomarkerKB provides a comprehensive and standardized representation of curated biomarkers, some limitations remain. For example, not all biomarker data sources explicitly report result changes (e.g., increased, decreased, presence, or absence), requiring manual inference in certain cases. This introduces a degree of variability in annotation despite extensive quality control and cross-verification measures. Furthermore, the current release of BiomarkerKB is predominantly centered on cancer-related biomarkers, reflecting the relative abundance of oncology data in structured public repositories. Broader coverage across additional disease domains is necessary and may be achievable through large-scale automated curation of unstructured text from scientific publications, electronic health records, clinical trial registries, drug labels, FDA reports, and related sources. While pipelines for integrating unstructured biomedical text, such as PubMed abstracts, clinical guidelines, and assay instructions for use, have been established by us in this work, the large-scale application and validation of large language models (LLMs) for automated extraction of biomarker-condition relationships is ongoing and an active area of research. These approaches will be critical for improving the depth, scalability, and automation of future data integration efforts. Future work will address these limitations through multiple complementary directions.

In conclusion, BiomarkerKB represents a foundational step toward a unified, scalable, and computationally enabled biomarker knowledgebase. By combining rigorous curation, ontology-driven standardization, and interactive web access, BiomarkerKB empowers researchers to explore, analyze, and discover biomarkers with improved reproducibility and translational relevance. Continued development will further extend its coverage, interoperability, and analytical capabilities, positioning BiomarkerKB as a core resource for precision medicine, systems biology, and biomarker-driven research. Furthermore, by organizing biomarker knowledge for seamless incorporation into interoperable knowledge graphs, BiomarkerKB establishes a foundation for integration with broader frameworks such as NSF Proto Open Knowledge Networks (Proto-OKN), an effort already initiated by our group. Such integration will allow biomarker data to participate in an increasingly interconnected and evolving landscape of scientific knowledge.

### License

BiomarkerKB is licensed under a permissive (CC BY 4.0) license. Users can modify the work and release their new version under any license they want (even a restrictive, commercial one), as long as they cite the resource.

## Acknowledgments

We would like to acknowledge the Biomarker Ontology Working Group (https://wiki.biomarkerkb.org/Biomaker_Ontology_Working_Group) for contributions and ideas for the controlled vocabulary and biomarker ontology that helped shape the biomarker data model.

This work has been funded by National Institute of Health award numbers U24OD038423 and OT2OD032092.

## Supplementary File Legends

Supplementary File 1. Biomarker Extraction using LLM, Agentic AI, and RAG

A) An LLM-driven, agentic workflow with MCP-backed tool orchestration. LOINC test codes are provided as input (1), and the AI agent processes them. If test descriptions are missing or incomplete, the agent retrieves them via automated LOINC web scraping (2). The enriched LOINC description is then analyzed by a large language model (LLM) to extract chemical (candidate biomarker)-condition pairs (3). The system evaluates whether biomarker extraction is conclusive (4). If conclusive, results are returned directly as structured output (9). If inconclusive (5), the AI agent triggers an evidence enrichment workflow by querying PubMed through either a Model Context Protocol (MCP) interface (6) or direct E-utilities access (7). Retrieved literature is indexed in a local SQLite full-text search (FTS5) database and accessed via retrieval-augmented generation (RAG) (8). The enriched evidence is then re-evaluated to resolve biomarker status, direction, and role, producing final structured outputs with explicit provenance. B) EDRN biomarkers are hosted in publicly available RDF files. These files contain free text with biomarker data. Trialed RAG methods to extract relevant text and then used LangChain model with Llama 3.3 to extract biomarker data. Biomarker data was then mapped to the biomarker data model C) OpenAI models were used to extract glycan biomarker data from PMC articles and abstracts. Glycan entities were identified and normalized. These were then paired with annotations and ontology IDs were mapped. This was then mapped to the biomarker data model for BiomarkerKB ingestion.

Supplementary File 2. Biomarker Entity Types Controlled Vocabulary File: biomarker_entity_types.txt

This file outlines and provides controlled entity terms for the biomarker entities that are present. The file provides definitions for the different entities, ontology ID and terms, and examples. This document is intended to be used with the other controlled vocabulary documents to provide standardized biomarker names.

Supplementary File 3. Biomarker Reporting Terms Controlled Vocabulary File: measured.txt

This file outlines and provides controlled entity terms for the reporting terms used in biomarker names. The file provides definitions for the different reporting terms (e.g., increase, decrease), ontology ID and terms, and examples. This document is intended to be used with the other controlled vocabulary documents to provide standardized biomarker names.

Supplementary File 4. Biomarker Aspect Controlled Vocabulary File: aspects.txt

This file outlines and provides controlled entity terms for the aspect terms used in biomarker names. The file provides definitions for the different aspects(e.g., level, expression), ontology ID and terms, and examples. This document is intended to be used with the other controlled vocabulary documents to provide standardized biomarker names.

Supplementary File 5. Biomarker Modifications Controlled Vocabulary File: modifications.txt

This file outlines and provides controlled entity terms for modifications that can be present in certain biomarkers. The file provides definitions for the different modifications (e.g., glycosylation, phosphorylation), ontology ID and terms, and examples. This document is intended to be used with the other controlled vocabulary documents to provide standardized biomarker names.

Supplementary File 6. Cypher Query for Increased Interleukin-6 biomarker knowledge graph construction.

This Cypher query example shows how to build the query to extract the nodes for increased interleukin-6 biomarker and connect them to breast carcinoma and known drug targets for this condition.

Supplementary File 7. Cypher Query for biomarker-centric network illustrating cross-disease and drug associations derived from breast cancer-associated genes.

This Cypher query example focuses on finding a different condition tied to different genes associated with breast cancer. This query also expands on finding relationships closest to the nearest degree to the nodes specified in the query.

Supplementary File 8. Cross-disease associations for druggable breast cancer-associated entities.

This table summarizes overlapping disease associations for breast cancer-associated genes with known drug interactions. For each gene, diseases outside the breast cancer hierarchy were retrieved through gene-disease association relationships. Relationship types reflect the underlying evidence connecting each gene to a disease. These results highlight potential shared molecular mechanisms across diverse disease contexts and demonstrate the utility of BiomarkerKB for identifying potential therapeutic repurposing opportunities and cross-disease relevance of clinically actionable entities.

Supplementary File 9. Cypher Query for Tankyrase Inhibitor Use Case Query

This Cypher query example shows how to build the query to extract the nodes for Tankyrase Inhibitor Use Case and connect to the associated target and drug nodes.

Supplementary File 10. Example box plot of annotations from aggregated electronic health record (EHR) data.

Box plot-based summaries of biomarker value distributions for cardiac troponin I (BiomarkerKB ID AN6373-1; UniProtKB accession 19429) and cardiac troponin T (BiomarkerKB ID AN6441-1; UniProtKB accession P54379) derived from EHR data. De-identified clinical EHR data were obtained from Oracle Health Real-World Data (Oracle Health Real World Data, 2025). Laboratory results identified using LOINC codes (https://loinc.org/) were mined and mapped to BiomarkerKB molecular entities. Named-entity recognition (NER) was performed using NextMove Leadmine (https://www.nextmovesoftware.com/leadmine.html) to map LOINC test descriptions to protein biomarkers. Unit harmonization followed by aggregation and stratification by age and sex was performed. A presumed-healthy cohort was defined using an inclusion and exclusion protocol informed by best practices in observational health data science, particularly those established by OHDSI (https://www.ohdsi.org/). Because identifying heathy population accurately is challenging a two-step process was applied, and the resulting box plots provide an overview of biomarker value distributions in presumed-healthy patients that potentially can be broadly applicable across diverse biomarkers. In the first step, to reduce the influence of noise and extreme values inherent in EHR-derived laboratory data, measurements were filtered to retain the middle 80% of the distribution centered on the median (https://github.com/unmtransinfo/cfde-biomarkers/). In the second step the box plots as shown in this figure and website were generated using the 1.5× interquartile range (IQR) criterion to identify and visually separate remaining extreme outliers, which may reflect subclinical or clinical conditions, assay-related effects, or pre-analytical variability. Furthermore, only biomarkers with median values within published normal reference ranges, as reported in the literature or authoritative sources, are displayed in BiomarkerKB; biomarkers with medians falling outside these ranges are flagged for further manual review. Together, these steps ensure that summarized reference distributions more accurately represent the central healthy population while highlighting entries that may warrant further investigation. In the specific case of troponin biomarkers, since the healthy population is defined as having troponin results below the 99th percentile, these data cannot be inferred to represent healthy cohorts but rather show basic distribution parameters (median, 1.5×IQR) varying among population cohorts. The goal is to provide users with a clinical data-centric, high-level overview that is driven by observed measurements rather than prior assumptions. This approach is intended to complement the published literature, not to replace it. Additional details are available for review.

