## Supplementary figures and images for "BiomarkerKB: An Integrated Knowledgebase Supporting Biomarker-Centric Exploration of Biomedical Data"

### Supplemental File 1

Supplementary File 1.

A

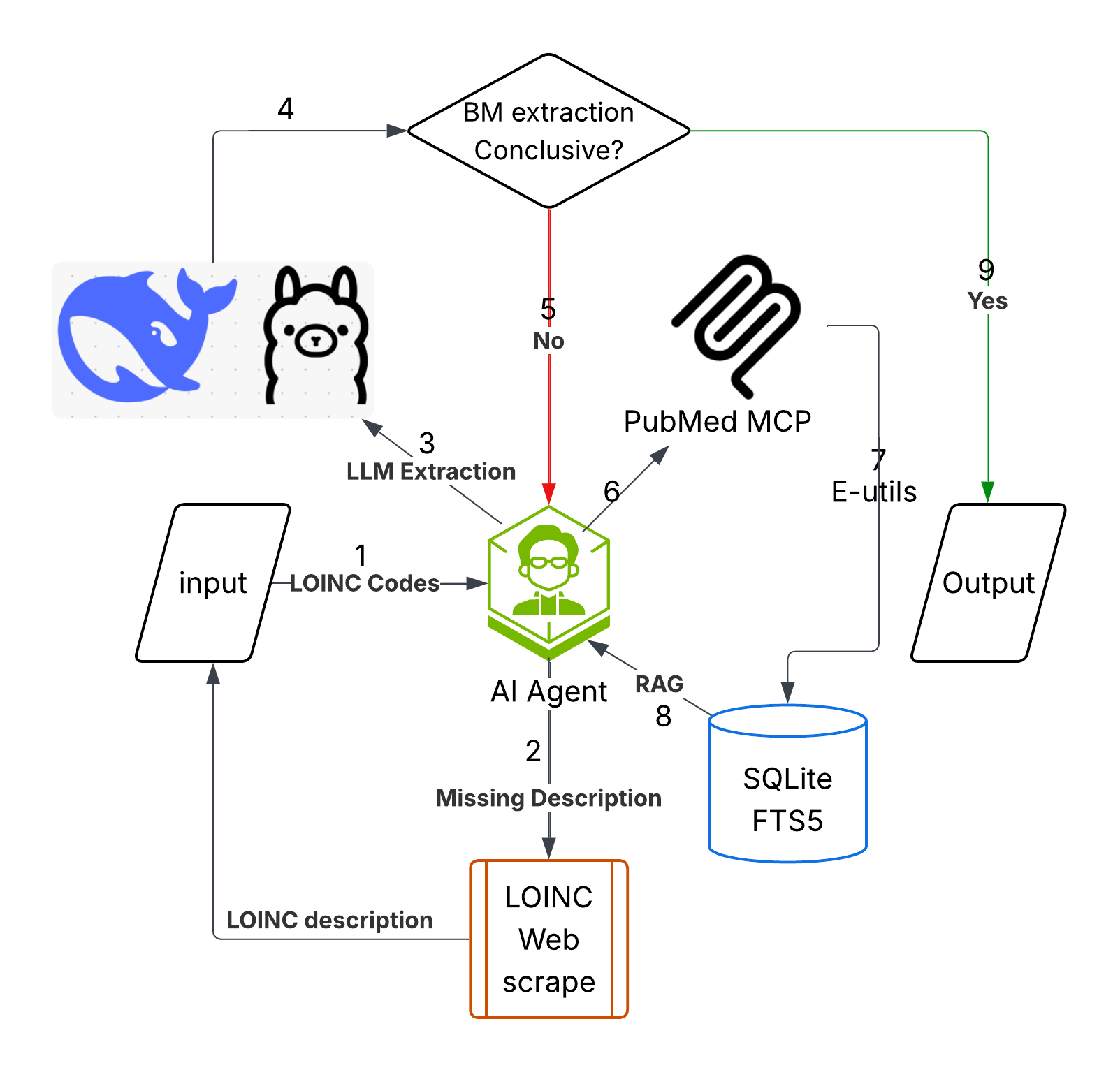

B

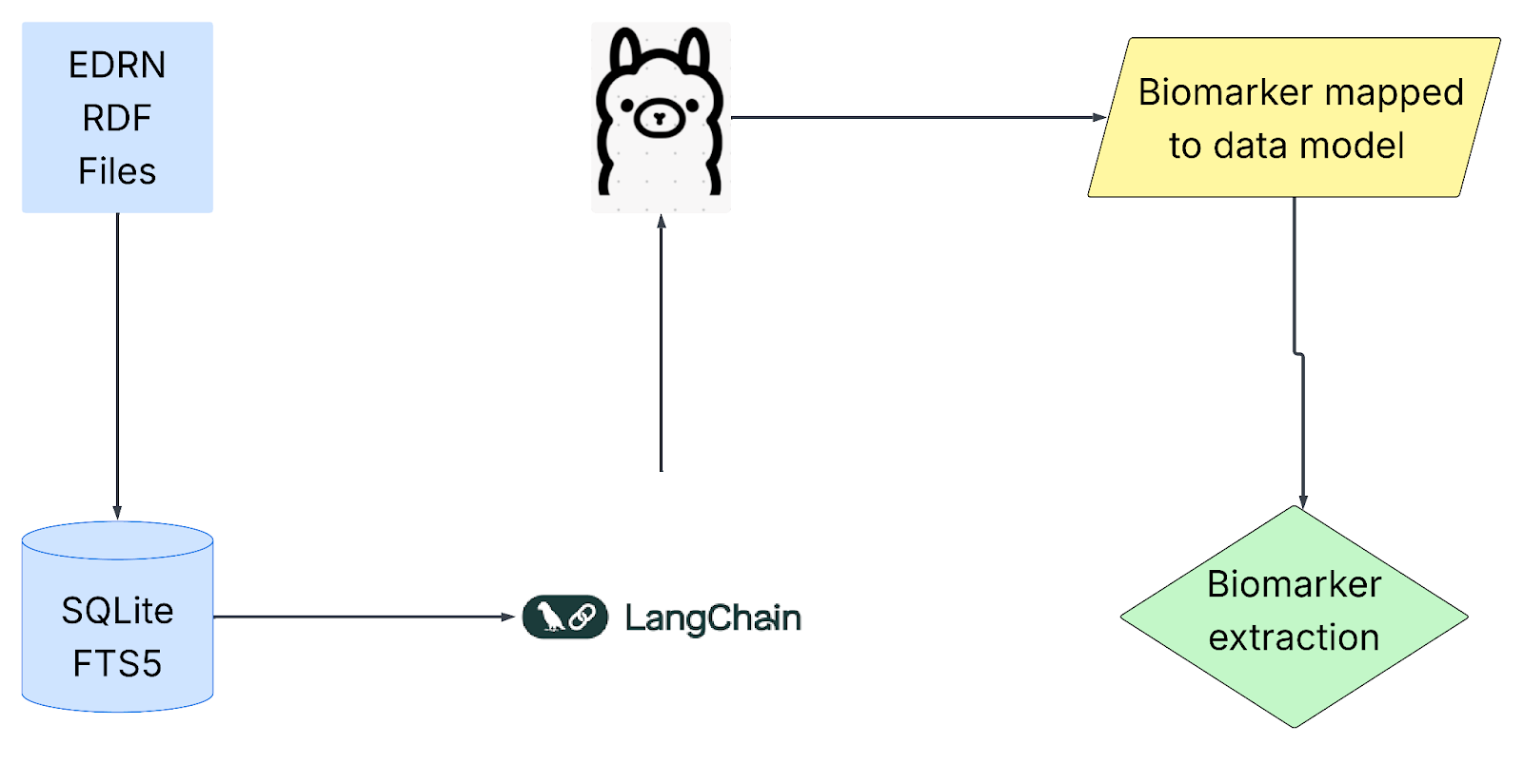

C

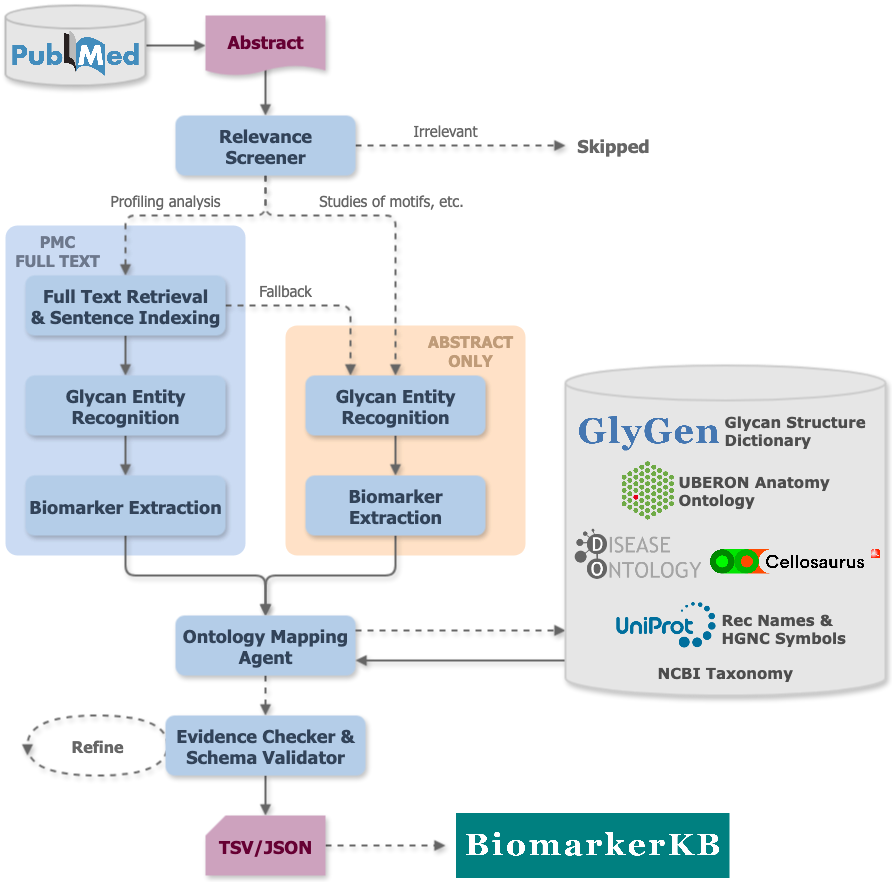

### Supplemental File 10

Supplementary File 10.Box plot of biomarker entity Troponin I

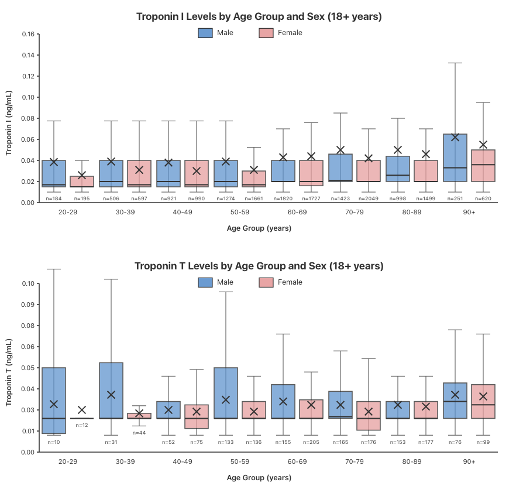
