## Supplemental File 2 for "BiomarkerKB: An Integrated Knowledgebase Supporting Biomarker-Centric Exploration of Biomedical Data"

Supplementary File 2.

```
--------------------------------------------------------------------------------
          BiomarkerKB Controlled Vocabulary
          Biomarker Entity Types Controlled Vocabulary File
          Biomarker Knowledgebase Project
--------------------------------------------------------------------------------

Description: Controlled vocabulary of biomarker entity types used in BiomarkerKB
Name:        biomarker_entity_types.txt
Release:     2025_07_16

--------------------------------------------------------------------------------

This document lists the biomarker entity types used in the BiomarkerKB
curation system. When a biomarker entity is paired with terms from
measurable.txt (increase, decrease, presence, absence, etc.), they form the
biomarker. The biomarker entity types are:

    * sequence variation
    * protein
    * metabolite
    * glycan
    * DNA
    * RNA
    * cell
    * lipid
    * image
    * mineral element

 Each entry consists of the following line codes:

  ----  ----------  ---------------------------  -------------------------------
  Code  Name        Content                      Occurrences
  ----  ----------  ---------------------------  -------------------------------
  ID    Identifier  Biomarker entity name        Once; starts an entry
  AC    Accession   Unique identifier (BM-xxxx)  Once
  DE    Definition  Definition/description       Once; line wrapping possible
  UF    Used For    Instructions for use         Once; line wrapping possible
  HI    Hierarchy                                Optional; zero or more
  SY    Synonym                                  Optional; zero or more
  EQ    Equivalent  External identifier with     Optional; zero or more
                      same meaning
  EX    Example     A biomarker that uses        Optional; zero or once; line
                      the indicated entity         wrapping possible
  NT    Notes                                    Optional; zero or once; line
                                                   wrapping possible
  //    Terminator  End of entry                 Once; ends an entry


--------------------------------------------------------------------------------

ID   sequence variation
AC   BM-0001
DE   For polymers, any difference between the observed order of monomer units
DE   within a polymer chain and the order within a suitably comparable reference
DE   polymer.
UF   Entities that differ when specifically comparing sequence; includes
UF   polymorphisms and mutations of genetic material and proteins.
HI   sequence variation
EQ   SO:0001059; sequence_alteration
EX   presence of rs11571833 mutation in BRCA2;
EX   https://biomarkerkb.org/biomarker/AN7363-3
//

ID   protein
AC   BM-0002
DE   Amino acid chain formed by ribosome-mediated translation of a genetically-
DE   encoded mRNA, and any post-translationally modified derivatives.
UF   Entities that are composed, in whole or in part, of protein; includes
UF   protein complexes and mixed-composition entities such as glycoproteins.
HI   protein
EQ   MESH:D011506; Proteins
EQ   PR:000000001; protein
EX   Interleukin-6; https://biomarkerkb.org/canonical/AN6278
NT   Mixed-composition entities can get multiple entity-type tags; for example,a
NT   glycoprotein would be both ‘protein’ and ‘glycan’.
//

ID   metabolite
AC   BM-0003
DE   A small molecule involved in metabolism.
UF   Entities that are inputs to or outputs from metabolic reactions.
HI   metabolite
EQ   CHEBI:25212; metabolite
EX   UREA; https://biomarkerkb.org/canonical/AN6341
NT   MESH term for metabolite being requested.
//

ID   glycan
AC   BM-0004
DE   Any oligosaccharide, polysaccharide or their derivatives consisting of
DE   monosaccharides or monosaccharide derivatives linked by glycosidic bonds.
UF   Entities that are composed, in whole or in part, of glycan; includes
UF   protein complexes and mixed-composition entities such as proteoglycans.
HI   glycan
EQ   CHEBI:167559
EX   N-glycan; https://biomarkerkb.org/canonical/AN6729
NT   Mixed-composition entities can get multiple entity-type tags; for example,a
NT   proteoglycan would be both ‘protein’ and ‘glycan’.
//

ID   DNA
AC   BM-0005
DE   A polymer of deoxyribose-containing nucleotides linked by phosphodiester
DE   bonds.
UF   Entities that are composed of DNA when measured in bulk, or that are direct
UF   or proxy measures of gene expression.
HI   DNA
EQ   MESH:D004247; DNA
EX   cfDNA; https://biomarkerkb.org/canonical/AN6380
//


ID   RNA
AC   BM-0006
DE   A polymer of ribose-containing nucleotides linked by phosphodiester bonds.
UF   Entities that are composed of RNA when measured in bulk, or that are direct
UF   or proxy measures of RNA expression.
HI   RNA
EQ   MESH:D012313; RNA
EX   miRNA-21; https://biomarkerkb.org/canonical/AN6498
//


ID   cell
AC   BM-0007
DE   An organism (or part thereof) that is a maximally connected compartment
DE   surrounded by a plasma membrane.
UF   Entities that are cells; includes cases when ratios between cell types are
UF   measured.
HI   cell
EQ   MESH:D002477; Cells
EQ   CL:0000000; cell
EX   WBC; https://biomarkerkb.org/canonical/AN6280
//

ID   lipid
AC   BM-0007
DE   A class of biomolecules including fats, oils, and certain hormones.
UF   Entities that are lipids.
HI   lipid
EQ   MESH:D008055; Lipids
EQ   CHEBI:18059; lipid
EX   very long chain fatty acid; https://biomarkerkb.org/canonical/AN6187
//

ID   image
AC   BM-0008
DE   Any visual display of structural or functional patterns of organs or tissues for
DE   clinical evaluation.
UF   Entities evaluated via the use of an image.
HI   image
EQ   NCIT:C48179; Image
EX   presence of ground-glass opacity;
EX   https://data.oncomx.org/allbiomarkers/biomarker/A0076
NT   For type ‘image’, the entity that is being imaged would be indicated as the
NT   sample source.
//

ID   mineral element
AC   BM-0009
DE   A chemical element occurring in biological systems, measured in elemental
DE   or ionic form (e.g., Na+, K+, Ca2+, Fe, Zn, Pb) as a nutrient, electrolyte,
DE   or toxicant.
UF   Entities that are elemental minerals or inorganic ions; includes electrolytes
UF   (Na+, K+, Cl−, Ca2+, Mg2+), trace elements (Fe, Zn, Cu, Se, I), and toxic
UF   metals/metalloids (Pb, Hg, Cd, As).
HI   mineral element
EQ
EX
//
```
