## Supplemental File 3 for "BiomarkerKB: An Integrated Knowledgebase Supporting Biomarker-Centric Exploration of Biomedical Data"

Supplementary File 3.

```
---------------------------------------------------------------------------
          BiomarkerKB Controlled Vocabulary:
          Biomarker Reporting Terms Controlled Vocabulary File
          Biomarker Knowledgebase Project
---------------------------------------------------------------------------

Description: Controlled vocabulary of standardized terms used to describe
biomarker status, behavior, or detection in curated data entries.
Name:        measured.txt
Release:     2025_07_16

---------------------------------------------------------------------------

  This document lists the controlled reporting terms used to describe
  biomarkers in the BiomarkerKB curation system. These nouns are used
  in fields like `biomarker` to standardize natural language descriptions.

  Each entry consists of the following line codes:
    ----  ----------  ---------------------------  ---------------------------------
  Code  Name        Content                      Occurrences
  ----  ----------  ---------------------------  ---------------------------------
  ID    Identifier  Reporting term               Once; starts an entry
  AC    Accession   Unique identifier (RT-xxxx)  Once
  DE    Definition  Definition/description       Once; line wrapping possible
  HI    Hierarchy                                Optional; zero or more
  SY    Synonym                                  Optional; zero or more
  EQ    Equivalent  External identifier with     Optional; zero or more
                      same meaning
  EX    Example                                  Optional; zero or once
  NT    Notes                                    Optional; zero or once; line
                                                   wrapping possible
  //    Terminator  End of entry                 Once; ends an entry

---------------------------------------------------------------------------

ID   increased
AC   RT-0001
DE   Indicates an assessed biomarker entity level is higher than normal by
DE   a clinically relevant degree.
HI   abundance
EQ   PATO:0002300; increased quality
EX   increased IL6 level; https://biomarkerkb.org/canonical/AN6278
//

ID   decreased
AC   RT-0002
DE   Indicates an assessed biomarker entity level is lower than normal by
DE   a clinically relevant degree.
HI   abundance
EQ   PATO:0002301; decreased quality
EX   decreased albumin level; https://biomarkerkb.org/canonical/AN6351
//

ID   presence of
AC   RT-0003
DE   Indicates that the assessed entity is present.
HI   presence
EX   presence of rs180177132 mutation in PALB2;
EX   https://biomarkerkb.org/canonical/AV9568
//


ID   difference
AC   RT-0004
DE   A statistic that is a subtraction of one quantity from another.
HI   expression
EQ   STATO:0000613; difference
EX   differential expression of TNMD; https://biomarkerkb.org/canonical/BB1486
//
```
