## Supplemental File 4 for "BiomarkerKB: An Integrated Knowledgebase Supporting Biomarker-Centric Exploration of Biomedical Data"

Supplementary File 4.

```
---------------------------------------------------------------------------
          BiomarkerKB Controlled Vocabulary:
          Biomarker Aspect Controlled Vocabulary File
          Biomarker Knowledgebase Project
---------------------------------------------------------------------------

Description: Controlled vocabulary of standardized aspects used to describe
the specific measurable or biological feature of a biomarker entity.
Name:        aspect.txt
Release:     2025_07_16
---------------------------------------------------------------------------

  This document lists controlled terms that represent measurable or
  definable aspects of a biomarker entity. These terms indicate what
  property, modification, or characteristic of the biomarker is being
  evaluated, enabling structured representation of natural language
  phrases such as "increased IL6 level" or "presence of BRCA2 mutation".

  Each entry consists of the following line codes:
    ----  ----------  ---------------------------  ---------------------------------
  Code  Name        Content                      Occurrences
  ----  ----------  ---------------------------  ---------------------------------
  ID    Identifier  Aspect term                  Once; starts an entry
  AC    Accession   Unique identifier (AS-xxxx)  Once
  DE    Definition  Definition/description       Once; line wrapping possible
  HI    Hierarchy                                Optional; zero or more
  EQ    Equivalent  External identifier with     Optional; zero or more
                      same meaning
  EX    Example                                  Optional; zero or once
  NT    Notes                                    Optional; zero or once; line
                                                   wrapping possible
  //    Terminator  End of entry                 Once; ends an entry
---------------------------------------------------------------------------

ID   level
AC   AS-0001
DE   A position on a scale measuring intensity, quality, or amount.
EQ   NCIT:C25554; Level
EX   increased IL6 level; https://biomarkerkb.org/canonical/AN6278
//

ID   expression
AC   AS-0002
DE   Production level or transcriptional activity of a gene or RNA molecule.
EQ   GO:0010467; gene expression
EX   differential expression of TNMD; https://biomarkerkb.org/canonical/BB1486
//

ID   mutation
AC   AS-0003
DE   A change in a sequence or structure in comparison to a reference entity
DE   due to an insertion, deletion or substitution event
DE   within a biomarker entity.
EQ   MI:0118; mutation
EX   presence of rs11571833 mutation in BRCA2;
EX   https://biomarkerkb.org/biomarker/AN7363-3
//
```
