## Supplemental File 5 for "BiomarkerKB: An Integrated Knowledgebase Supporting Biomarker-Centric Exploration of Biomedical Data"

Supplementary File 5.

```
---------------------------------------------------------------------------
          BiomarkerKB Controlled Vocabulary:
          Biomarker Modifications Controlled Vocabulary File
          Biomarker Knowledgebase Project
---------------------------------------------------------------------------

Description: Controlled vocabulary of molecular modifications representing
chemical or post-translational changes to biomarker entities.
Name:        modifications.txt
Release:     2025_07_16
---------------------------------------------------------------------------

  This document lists controlled modification terms that describe chemical,
  structural, or enzymatic alterations to biomarker entities. These terms
  are used to represent states such as methylation, phosphorylation, or
  glycosylation, enabling consistent annotation of modified biomarker forms.

  Each entry consists of the following line codes:
    ----  ----------  ---------------------------  ---------------------------------
  Code  Name        Content                      Occurrences
  ----  ----------  ---------------------------  ---------------------------------
  ID    Identifier  Modification term            Once; starts an entry
  AC    Accession   Unique identifier (MO-xxxx)  Once
  DE    Definition  Definition/description       Once; line wrapping possible
  HI    Hierarchy                                Optional; zero or more
  EQ    Equivalent  External identifier with     Optional; zero or more
                      same meaning
  EX    Example                                  Optional; zero or once
  NT    Notes                                    Optional; zero or once; line
                                                   wrapping possible
  //    Terminator  End of entry                 Once; ends an entry
---------------------------------------------------------------------------

ID   methylation
AC   MO-0001
DE   The process in which a methyl group is covalently attached to a molecule.
HI   chemical modification
EQ   GO:0032259; methylation
EX   presence of methylation in DCC; https://biomarkerkb.org/canonical/AN6479
//


ID   phosphorylation
AC   MO-0002
DE   The process of introducing a phosphate group into a molecule, usually
DE   with the formation of a phosphoric ester, a phosphoric anhydride or a
DE   phosphoric amide.
HI   post-translational modification
EQ   GO:0016310; phosphorylation
EX   AKT1 S473 Phosphorylation; https://biomarkerkb.org/canonical/AN4659
//


ID   acetylation
AC   MO-0003
DE   Acetylation involves the covalent linkage of an acetyl group into an
DE   organic molecule.
HI   post-translational modification
EQ   NCIT:C16255; Acetylation
//


ID   glycosylation
AC   MO-0004
DE   The covalent chemical or biochemical addition of carbohydrate or glycosyl
DE   groups to other chemicals, by glycosyl transferases.
HI   post-translational modification
EQ   NCIT:C21034; Glycosylation
//
```
