## Supplemental File 6 for "BiomarkerKB: An Integrated Knowledgebase Supporting Biomarker-Centric Exploration of Biomedical Data"

Supplementary File 6

// IL-6 → BIOMARKER:AN6278-4 CUI → Breast Carcinoma → PubChem compounds

WITH ['interleukin-6','interleukin 6','il6','il-6'] AS names, 'Breast Carcinoma' AS TARGET

// Resolve IL-6 by name

MATCH (il6:Concept)-[:PREF_TERM]->(ilt:Term)

WHERE toLower(ilt.name) IN names OR toLower(ilt.name) CONTAINS 'interleukin-6'

// IL-6 ↔ biomarker (use type filter to avoid the :A|:B issue)

MATCH p_il6_bio = (il6)-[rb]-(b:Concept)

WHERE type(rb) IN ['indicated_by_above_normal_level_of','inverse_indicated_by_above_normal_level_of']

AND b.CUI = 'BIOMARKER:AN6278-4 CUI'

// biomarker → condition (keep intermediate node `x` so edges render)

MATCH p_bio_cond = (b)-[r1]-(x:Concept)-[r2]-(cond:Concept)-[:PREF_TERM]->(ct:Term)

WHERE toLower(ct.name) = toLower(TARGET)

// condition → compounds (PubChem) as paths

OPTIONAL MATCH p_cond_cmp = (cond)-[rc]-(cmp:Concept)-[:CODE]->(:Code {SAB:'PUBCHEM'})

// ---- add PREF_TERM paths for every Concept we’re returning ----

WITH p_il6_bio, p_bio_cond, p_cond_cmp,

collect(DISTINCT il6)+collect(DISTINCT b)+collect(DISTINCT x)+collect(DISTINCT cond)+

[cmp IN nodes(p_cond_cmp) WHERE cmp:Concept] AS concept_lists

WITH p_il6_bio, p_bio_cond, p_cond_cmp,

apoc.coll.flatten(concept_lists) AS concepts

UNWIND concepts AS n

OPTIONAL MATCH p_term = (n)-[:PREF_TERM]->(t:Term)

// Return all paths so the Browser can draw everything connected

RETURN DISTINCT p_il6_bio, p_bio_cond, p_cond_cmp, p_term;
