## Supplemental File 7 for "BiomarkerKB: An Integrated Knowledgebase Supporting Biomarker-Centric Exploration of Biomedical Data"

Supplementary File 7

// VISUALIZE: BrCa SNOMED root -> top druggable biomarkers -> top overlapping diseases

// Uses APOC virtual relationships so it renders cleanly in Neo4j Browser

// Caps: top 5 biomarkers, top 5 overlap diseases per biomarker

// Requires APOC

WITH '126926005' AS snomedCode

// --- root + SNOMED descendant subtree ---

MATCH (sn:Code)

WHERE sn.SAB STARTS WITH 'SNOMED' AND toString(sn.CODE) = snomedCode

MATCH (root:Concept)-[:CODE]-(sn)

MATCH p = (root)-[:inverse_isa*0..10]->(bc:Concept)

WHERE all(r IN relationships(p) WHERE coalesce(r.SAB,'') STARTS WITH 'SNOMED')

WITH root, sn, collect(DISTINCT bc) AS bcConcepts

WITH root, sn, bcConcepts, apoc.coll.toSet([x IN bcConcepts | elementId(x)]) AS bcEids

// --- biomarkers (genes) connected to the BrCa subtree ---

UNWIND bcConcepts AS bc

MATCH (bc)-[gd]-(bio:Concept)

WHERE type(gd) IN [

'disease_has_associated_gene',

'gene_associated_with_disease',

'gene_associated_with_disease_or_phenotype',

'inverse_gene_associated_with_disease_or_phenotype',

'gene_product_malfunction_associated_with_disease',

'associated_with_malfunction_of_gene_product'

]

WITH root, sn, bcEids, bio, count(DISTINCT bc) AS supportBC

// --- druggability count (gene -> drug OR gene -> protein -> drug) ---

CALL {

WITH bio

CALL {

WITH bio

MATCH (bio)-[:has_target|is_target|ro]-(drug:Concept)

WHERE EXISTS { MATCH (drug)-[:CODE]->(rx:Code) WHERE rx.SAB STARTS WITH 'RXNORM' }

RETURN collect(DISTINCT drug) AS ds

UNION

WITH bio

MATCH (bio)-[:gene_encodes_gene_product|gene_product_encoded_by_gene]-(gp:Concept)

MATCH (gp)-[:has_target|is_target|ro]-(drug:Concept)

WHERE EXISTS { MATCH (drug)-[:CODE]->(rx:Code) WHERE rx.SAB STARTS WITH 'RXNORM' }

RETURN collect(DISTINCT drug) AS ds

}

WITH apoc.coll.toSet(apoc.coll.flatten(collect(ds))) AS drugs

RETURN size(drugs) AS drugCount

}

WITH root, sn, bcEids, bio, supportBC, drugCount

WHERE drugCount > 0

WITH root, sn, bcEids, bio, supportBC, drugCount

ORDER BY drugCount DESC, supportBC DESC

LIMIT 5

// Make a display node so drugCount shows in the graph label

CALL apoc.create.vNode(

['BIO_DISPLAY'],

{

CUI: bio.CUI,

prefName: coalesce(bio.prefName,'') + ' (drugs=' + toString(drugCount) + ', BC=' + toString(supportBC) + ')',

baseName: coalesce(bio.prefName,''),

drugCount: drugCount,

supportBC: supportBC

}

) YIELD node AS bioDisplay

// Root -> biomarker edge (virtual)

CALL apoc.create.vRelationship(

root,

'TOP_DRUGGABLE_BIOMARKER',

{edgeLabel:'top druggable biomarkers'},

bioDisplay

) YIELD rel AS rBio

// --- overlap diseases outside the BrCa subtree ---

MATCH (bio)-[r]-(d:Concept)

WHERE type(r) IN [

'disease_has_associated_gene',

'gene_associated_with_disease',

'gene_associated_with_disease_or_phenotype',

'inverse_gene_associated_with_disease_or_phenotype'

]

AND d <> root

AND NOT elementId(d) IN bcEids

AND coalesce(d.prefName,'') <> ''

AND NOT d.prefName IN ['Neoplasms','Malignant Neoplasms','Benign Neoplasm']

// ✅ keep rBio in scope from here onward

WITH root, bioDisplay, rBio, d, collect(DISTINCT type(r)) AS relTypes

ORDER BY d.prefName

WITH root, bioDisplay, rBio, collect({d:d, relTypes: relTypes})[..5] AS overlaps

UNWIND overlaps AS ov

WITH root, bioDisplay, rBio, ov.d AS d, ov.relTypes AS relTypes

CALL apoc.create.vRelationship(

bioDisplay,

'UNEXPECTED_DISEASE_OVERLAP',

{edgeLabel: apoc.text.join(relTypes,' | ')},

d

) YIELD rel AS rOverlap

RETURN root, bioDisplay, d, rBio, rOverlap;
