## Supplemental File 8 for "BiomarkerKB: An Integrated Knowledgebase Supporting Biomarker-Centric Exploration of Biomedical Data"

Supplementary File 8.

| **Gene** | **GeneCUI** | **Support (BC)** | **Drug Count** | **Overlapping Disease** | **Disease CUI** | **Relationship Types** |
| --- | --- | --- | --- | --- | --- | --- |
| BRAF gene | C0812241 | 1 | 7 | Acrania | C0702169 | gene_associated_with_disease_or_phenotype; inverse_gene_associated_with_disease_or_phenotype |
| BRAF gene | C0812241 | 1 | 7 | Adenocarcinoma of lung | C0152013 | gene_associated_with_disease_or_phenotype; inverse_gene_associated_with_disease_or_phenotype |
| BRAF gene | C0812241 | 1 | 7 | Adenocarcinoma of prostate | C0007112 | gene_associated_with_disease_or_phenotype; inverse_gene_associated_with_disease_or_phenotype |
| ERBB2 gene | C0242957 | 1 | 14 | Gastric adenocarcinoma | C0278701 | gene_associated_with_disease_or_phenotype; inverse_gene_associated_with_disease_or_phenotype |
| ERBB2 gene | C0242957 | 1 | 14 | Colorectal carcinoma | C0009402 | gene_associated_with_disease_or_phenotype; inverse_gene_associated_with_disease_or_phenotype |
| ERBB2 gene | C0242957 | 1 | 14 | Adenocarcinoma of lung | C0152013 | gene_associated_with_disease; disease_has_associated_gene; gene_associated_with_disease_or_phenotype |
| ALK gene | C1332080 | 1 | 9 | Neuroblastoma | C0027819 | gene_associated_with_disease_or_phenotype; inverse_gene_associated_with_disease_or_phenotype |
| ALK gene | C1332080 | 1 | 9 | Brain neoplasms | C0006118 | gene_associated_with_disease_or_phenotype; inverse_gene_associated_with_disease_or_phenotype |
| IDH1 gene | C1415876 | 1 | 3 | Glioblastoma multiforme | C1621958 | gene_associated_with_disease_or_phenotype; inverse_gene_associated_with_disease_or_phenotype |
| MTOR gene | C1414805 | 1 | 3 | Cutaneous melanoma | C0151779 | gene_associated_with_disease_or_phenotype; inverse_gene_associated_with_disease_or_phenotype |
