## Supplemental File 9 for "BiomarkerKB: An Integrated Knowledgebase Supporting Biomarker-Centric Exploration of Biomedical Data"

Supplementary File 9. Tankyrase Inhibitor Use Case Query

WITH "C5418065" AS rootCUI

// 1) Root concept (center purple node)

MATCH (root:Concept {CUI: rootCUI})

// 2) Pull the neighborhood seen in the figure:

//	- CODE links (to Code nodes like C174060, C21235, etc.)

//	- Semantic type links (STY) to type/category nodes

//	- “subset/includes” style edges between Concept nodes

//	- Drug-target links (has_target / is_target / ro)

//	- Gene/protein encoding links

OPTIONAL MATCH (root)-[rCode:CODE]->(rootCode:Code)

OPTIONAL MATCH (root)-[rSty:STY]->(sty:Concept)

OPTIONAL MATCH (root)-[rSubset:concept_in_subset|subset_includes_concept|inverse_concept_in_subset|inverse_subset_includes_concept]-(nearConcept:Concept)

// Targets (genes/proteins) directly linked from the root drug/chemical

OPTIONAL MATCH (root)-[rTarget:has_target|is_target|ro]-(target:Concept)

// If the target is a gene/protein product, try to traverse to the corresponding gene (like your "Tankyrase Gene"/"TNKS Gene")

OPTIONAL MATCH (target)-[rEnc:gene_product_encoded_by_gene|gene_encodes_gene_product]-(gene:Concept)

// Codes/synonyms for the gene side (CODE / PT / SY are shown in your figure)

OPTIONAL MATCH (gene)-[rGeneCode:CODE]->(geneCode:Code)

OPTIONAL MATCH (gene)-[rGenePT:PT]->(genePT:Concept)

OPTIONAL MATCH (gene)-[rGeneSY:SY]->(geneSY:Concept)

// Also show any CODE / PT / SY on the target node itself (sometimes targets are already the “gene/protein” concept)

OPTIONAL MATCH (target)-[rTCode:CODE]->(tCode:Code)

OPTIONAL MATCH (target)-[rTPT:PT]->(tPT:Concept)

OPTIONAL MATCH (target)-[rTSY:SY]->(tSY:Concept)

// 3) Return everything as a graph

RETURN

root,

rootCode, rCode,

sty, rSty,

nearConcept, rSubset,

target, rTarget,

gene, rEnc,

geneCode, rGeneCode,

genePT, rGenePT,

geneSY, rGeneSY,

tCode, rTCode,

tPT, rTPT,

tSY, rTSY;
